# Ecological network assembly: how the regional metaweb influences local food webs

**DOI:** 10.1101/340430

**Authors:** Leonardo A. Saravia, Tomás I. Marina, Nadiah P. Kristensen, Marleen De Troch, Fernando R. Momo

**Affiliations:** Instituto de Ciencias, Universidad Nacional de General Sarmiento, J.M. Gutierrez 1159 (1613), Los Polvorines, Buenos Aires, Argentina; INEDES, Universidad Nacional de Luján, CC 221, 6700 Luján, Argentina; Centro Austral de Investigaciones Científicas (CADIC-CONICET), Ushuaia, Argentina; Marine Biology, Ghent University, Krijgslaan 281/S8, B-9000, Ghent, Belgium; Department of Biological Sciences, National University of Singapore, 14 Science Drive 4 Singapore 117543, Singapore

**Keywords:** Metaweb, ecological network assembly, network assembly model, food web structure, modularity, trophic coherence, motif, topological roles, null models

## Abstract

1. Local food webs result from a sequence of colonisations and extinctions by species from the regional pool or metaweb, i.e., the assembly process. Assembly is theorised to be a selective process: whether or not certain species or network structures can persist is partly determined by local processes including habitat filtering and dynamical constraints. Consequently, local food web structure should reflect these processes.
2. The goal of this study was to test evidence for these selective processes by comparing the structural properties of real food webs to the expected distribution given the metaweb. We were particularly interested in ecological dynamics; if the network properties commonly associated with dynamical stability are indeed the result of stability constraints, then they should deviate from expectation in the direction predicted by theory.
3. To create a null expectation, we used the novel approach of randomly assembling model webs by drawing species and interactions from the empirical metaweb. The assembly model permitted colonisation and extinction, and required a consumer species to have at least one prey, but had no habitat type nor population dynamical constraints. Three data sets were used: (1) the marine Antarctic metaweb, with 2 local food-webs; (2) the 50 lakes of the Adirondacks; and (3) the arthropod community from Florida Keys’ classic defaunation experiment.
4. Contrary to our expectations, we found that there were almost no differences between empirical webs and those resulting from the null assembly model. Few empirical food webs showed significant differences with network properties, motif representations and topological roles. Network properties associated with stability did not deviate from expectation in the direction predicted by theory.
5. Our results suggest that — for the commonly used metrics we considered — local food web structure is not strongly influenced by dynamical nor habitat restrictions. Instead, the structure is inherited from the metaweb. This suggests that the network properties typically attributed as causes or consequences of ecological stability are instead a by-product of the assembly process (i.e., spandrels), and may potentially be too coarse to detect the true signal of dynamical constraint.

## Introduction

What determines the structure of a food web? The characterization of ecological systems as networks of interacting elements has a long history (Paine, 1966; May, 1972; Cohen & Newman, 1985); however, the effects of ecological dynamical processes on network structure are not fully understood. Structure is the result of community assembly, which is a repeated process of species arrival, colonization, and local extinction (Cornell & Harrison, 2014). That implies there are two major components that determine food web structure: the composition of the regional pool, from which the species are drawn; and a selective process, which determines which species can arrive and persist in the local web. The selective process is very complex and involves multiple mechanisms (Mittelbach & Schemske, 2015). However, we should in theory be able to detect the signal of this process by comparing local webs to the regional pool from which they were drawn.

The structure of a food web is ultimately constrained by the species and potential interactions that exist in the regional pool, i.e., the metaweb. The regional pool is shaped by evolutionary and biogeographical processes that imply large spatial and temporal scales (Carstensen, Lessard, Holt, Krabbe Borregaard, & Rahbek, 2013; Kortsch et al., 2018), and it generally extends over many square kilometers and contains a large number of habitats and communities (Mittelbach & Schemske, 2015). Each of the local communities that make up the metaweb can have different food web structures, both in terms of the species present and interactions between them. Consequently, the metaweb includes many species co-occurrence and interaction possibilities that do not occur in reality.

Within the ultimate constraint imposed by the metaweb, the composition of the local community is determined by metacommunity processes. Which regional species can arrive and persist in a web is influenced by dispersal, environmental filters, biotic interactions and stochastic events (HilleRisLambers, Adler, Harpole, Levine, & Mayfield, 2012). These processes have been studied using metacommunity theory, where different spatial assemblages are connected through species dispersal (Leibold, Chase, & Ernest, 2017). Recently, there has been an increase in food web assembly studies, integrating them with island biogeography (Gravel, Massol, Canard, Mouillot, & Mouquet, 2011), metacommunity dynamics (Pillai, Gonzalez, & Loreau, 2011; Liao, Chen, Ying, Hiebeler, & Nijs, 2016) and the effects of habitat fragmentation (Mougi & Kondoh, 2016). As an extension of the species-area relationship (SAR) approach, one can derive a network-area relationship (NAR) using theoretical models (Galiana et al., 2018). However, this approach assumes that ecological dynamics (e.g., stability) will have no influence. Compared to the body of metacommunity theory, there are very few studies that have analyzed the assembly process using experimental or empirical data, and none of them focuses on topological network properties that could be related to different assembly processes. Piechnik, Lawler, & Martinez (2008) found that the first to colonize are trophic generalists followed by specialists, supporting the hypothesis that biotic interactions are important in the assembly process (Holt, Lawton, Polis, & Martinez, 1999). Baiser, Buckley, Gotelli, & Ellison (2013) showed that habitat characteristics and dispersal capabilities were the main drivers of the assembly. Fahimipour & Hein (2014) also found that colonization rates were an important factor.

On top of metacommunity processes, local dynamical processes play a role in determining food web structure, and the potential for stability to constrain food-web structure has received plenty of theoretical attention (May, 1972; K. S. McCann, 2000). Some theorists conceive of assembly as a non-Darwinian selection process (Borrelli, 2015), whereby species and structures that destabilize the web will be lost and stabilizing structures persist (Pawar, 2009; Borrelli, 2015). Typically, assembly simulations produce large webs that are both stable in the dynamical sense and relatively resistant to further invasions (Drake, 1990; Luh & Pimm, 1993; Law & Morton, 1996). Therefore, we expect that particular structural properties that confer stability will be over-represented in real food webs (Borrelli et al., 2015), as these are the webs that are able to persist in time (Grimm, Schmidt, & Wissel, 1992).

There is some evidence that real food webs possess stabilizing structural properties. A classic finding is that dynamical models parameterized with realistic species interaction-strength patterns have higher stability than randomized alternatives (de Ruiter, Neutel, & Moore, 1995; Neutel, Heesterbeek, & Ruiter, 2002). The frequency of 3-node sub-networks, called motifs (Milo et al., 2002), is correlated with the stability in ecological (Borrelli, 2015) and other biological (Prill, Iglesias, & Levchenko, 2005) networks. However, stability-enhancing structural features can also arise for non-dynamical reasons. For example, the nested structure of mutualistic networks can arise as a spandrel of adaptive radiation (Maynard, Serván, & Allesina, 2018; Valverde et al., 2018), and low connectance may occur as a consequence of restricted diet breadth and adaptive foraging behavior (Beckerman, Petchey, & Warren, 2006). Furthermore, it is niche models — which are typically interpreted in terms of physiological constraints on predation relationships and do not rely upon population dynamic mechanisms— that have been most successful at reproducing realistic food web structure (R. J. Williams & Martinez, 2000; Loeuille & Loreau, 2005). This raises the possibility that the structural attributes typically measured in real webs can be explained by other processes, or may be too coarse to detect a subordinate influence of dynamics.

To test the hypothesis that dynamical selective processes are responsible for food web structure, we need an appropriate null model (Lau, Borrett, Baiser, Gotelli, & Ellison, 2017). Here, we propose that the metaweb itself can be used to create that baseline for comparison. We conceive of the metaweb as the source of foodweb structural diversity, from which local food web structure is drawn, and upon which local processes can act. Although the metaweb is also a consequence of local assembly processes (being the sum of local webs), it contains species co-occurrences and network structures that never occur in a local web, including those presumably precluded by local dynamics. Therefore, if there are local selective processes that determine the structure of local food webs, then comparing local webs to the metaweb may allow us to separate the larger evolutionary and biogeographical processes from the theorized local selective process. For example, we would expect to find that the structural properties that confer stability are over-represented in local food webs compared to the metaweb.

In this study, we developed a null model independent of dynamic stability processes and compared the resulting structure to real food webs using network properties. We cannot directly compare the metaweb properties with the local webs properties since they are dependent on size, number of links and/or connectance (Dunne, Williams, & Martinez, 2002; Poisot & Gravel, 2014). Therefore, we compared the real networks to networks generated by the null model, which takes into account this issue. To create null food webs, we made the most minimal assumption possible about the metacommunity process: any species from the metaweb can colonize and persist in a local web given at least one prey (food) item available. Thus the model considers colonization-extinction and secondary extinctions events constrained by network structure, so it does not include dynamical stability and local habitats constraints that are thought to drive the assembly process. If the real food web structure differs from null models and in the direction predicted by theory, then that is evidence in favour of the hypothesis that food web structure is constrained by dynamics.

## Methods

We compiled three metawebs with their corresponding local food webs, with a total of 58 local food webs from a variety of regions and ecosystems. We built the first metaweb from the Southern Ocean database compiled by Raymond et al. (2011), selecting only species located at latitudes higher than 60°S. Raymond et al. (2011) compiled information from direct sampling methods of dietary assessment, including gut, scat, and bolus content analysis, stomach flushing, and observed feeding. We considered that the metaweb is the regional pool of species defined by the biogeographic Antarctic region. As local food webs we included two of the most well-resolved datasets publicly available for the region: Weddell Sea and Potter Cove food webs. The first includes species situated between 74°S and 78°S with a West-East extension of approximately 450 km and comprises all information about trophic interactions available for the zone since 1983 (Jacob et al., 2011), this dataset was obtained from Brose et al. (2005). The Potter Cove food web comes from a 4 km long and 2.5 km wide Antarctic fjord located at 62°14’S, 58°40’W, South Shetland Islands (Marina et al., 2018). To make datasets compatible, we firstly checked taxonomic names for synonyms, and secondly added species (either prey or predator) with their interactions to the metaweb when the local food webs contain a greater taxonomic resolution. This resulted in the addition of 258 species to the metaweb, which represent 33% of the total. We named this the Antarctic metaweb, which has 846 species (*S*), 6897 links (*L*) and a connectance (*L*/ *S*^2^) of 0.01.

The second metaweb was collected from pelagic organisms of 50 lakes of the Adirondacks region (Havens, 1992), which were sampled once during summer 1984 (Sutherland, 1989). Havens (1992) determined the potential predator–prey interactions among 211 species from previous diet studies; species that lacked a trophic link were deleted and feeding links were assumed when the species involved were present in a particular lake. The so-called Lakes metaweb considers 211 species, 8426 links and a connectance of 0.19, this was obtained from the GATEWAy database (Brose et al., 2019).

The third metaweb comes from a well-known defaunation experiment performed in the Florida Keys in the 1960’s (Simberloff & Wilson, 1969; Piechnik, Lawler, & Martinez, 2008), where six islands of 11–25 meters in diameter were defaunated with insecticide. The arthropods were censused before the experiment and after it approximately once every 3 weeks during the first year and again 2 years after defaunation. For the metaweb and local webs we used only the first census that represent a complete community. Piechnik, Lawler, & Martinez (2008) determined the trophic interactions among 155 species (5114 links, connectance 0.21) using published information and expert opinions. This dataset was obtained directly from the authors of Gravel, Massol, Canard, Mouillot, & Mouquet (2011).

### Metaweb Assembly Null Model

To consider network assembly mechanisms we used a metaweb assembly model (Figure 1), similar to the trophic theory of island biogeography (Gravel, Massol, Canard, Mouillot, & Mouquet, 2011). In this model species migrate from the metaweb to a local web with a probability *c*, and become extinct from the local web with probability *e*; a reminiscence of the theory of island biogeography (MacArthur & Wilson, 1967), but with the addition of network structure. Species migrate with their potential network links from the metaweb, then in the local web, species have a probability of secondary extinction *se* if none of its preys are present, which only applies to non-basal species. When a species goes extinct locally it may produce secondary extinctions modulated by *se* (Figure 1).

**Figure 1:**
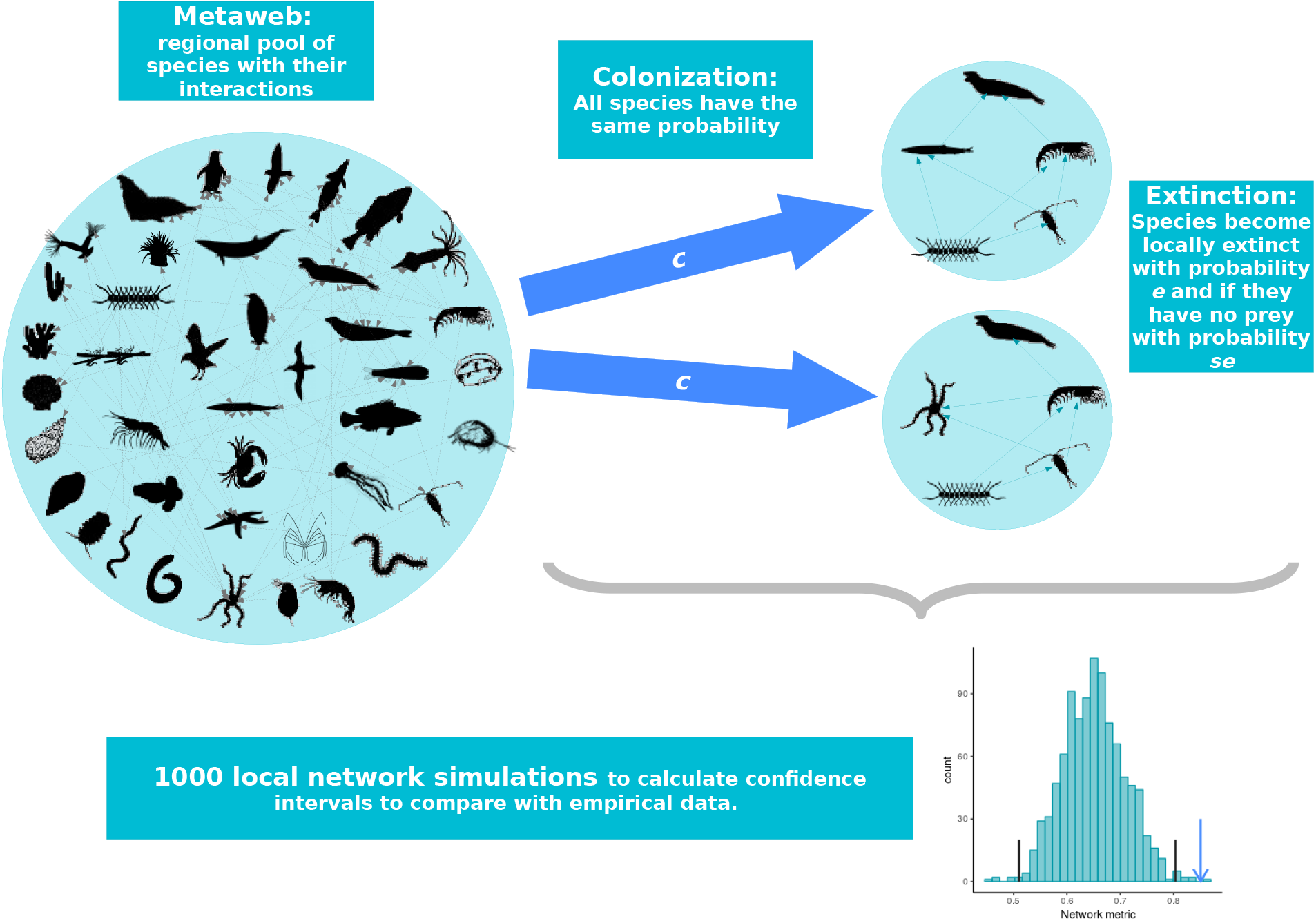
Schematic diagram of the metaweb assembly null model: species migrate from the metaweb with a probability *c* to a local network carrying their potential links; here they have a probability of extinction *e*. Additionally, predators become extinct if their preys are locally extinct with probability *se*. We simulate 1000 local networks and calculate network properties. From the distribution of these topological properties we calculate 1% confidence intervals to compare with empirical networks

Then there are three possible events: colonization, extinction, and secondary extinction. After a colonization event with probability *c*, the species is present in the local community and two other events are possible:

1. if it is a basal species it does not need predators to survive, then it persists until an extinction event with probability *e*;
2. if it is a non-basal species it could become extinct with probability *e* but if it has no prey it could also become extinct with probability *se*.

These events could happen at random if the necessary conditions are fulfilled, to simulate the model we use the Gillespie (1976) algorithm that produces an statistically exact trajectory of the stochastic process (Black & McKane, 2012).

We simulated this model in time and it eventually reached a steady state that depends on the migration and extinction probabilities but also on the structure of the metaweb. The ratio of immigration vs. extinction *α* = *c*/*e* is hypothesized to be inversely related to the distance to the mainland (MacArthur & Wilson, 1967), and as extinction should be inversely proportional to population size (Hanski, 1999), the ratio *α* is also hypothesized to be related to the local area.

For the model used in Gravel, Massol, Canard, Mouillot, & Mouquet (2011), simulations with the same ratio *α* = *c*/*e* should give the same results, but as our model incorporates *se* as an additional parameter this might not be the case. We checked this performing simulations with different combinations of *c, e*, and *se* keeping *α* constant for different metawebs. We found differences for some of the combinations (Figure S6), thus we performed the fitting using the 3 parameters.

To fit the model to each metaweb we performed 150000 simulations with a wide range of parameters (Table S1) using latin hypercube sampling (Fang, Li, & Sudjianto, 2005). We simulated the model for 1000 time steps and use the last 100 time steps to calculate averages for the number of species *S*_*m*_, the number of links *E*_*m*_ and the connectance 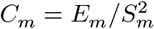. Then we calculated a relative distance to the number of species *S*_*e*_ and connectance *C*_*e*_ of the empirical food webs:

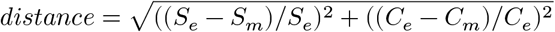

Then we used the parameters with the minimal distance to simulate the model and compare with the network properties described in the following section. The fitted parameters for all local food webs are presented in table S2.

In summary, this model considers colonization-extinction and secondary extinctions events constrained by network structure, with no consideration of population dynamics and interaction strength. Then, this simple model acts as a null model: if we observe a deviation from a network property obtained with the null model then those mechanisms that are excluded from the model may be acting (de Bello, 2012).

### Structural network properties

We first calculated trophic coherence (Johnson, Domínguez-García, Donetti, & Muñoz, 2014), that is related to stability in the sense that small perturbations could get amplified or vanished, which is called local linear stability (May, 1972; Rohr, Saavedra, & Bascompte, 2014). A food web is more coherent when *Q* is closer to zero, thus the maximal coherence is achieved when *Q* = 0, and corresponds to a layered network in which every node has an integer trophic level (Johnson, Domínguez-García, Donetti, & Muñoz, 2014; Johnson & Jones, 2017). A related metric is mean trophic level, historically used as an ecosystem health indicator (Pauly, Christensen, Dalsgaard, Froese, & Torres, 1998), predicting that food webs with higher trophic levels are less stable (Borrelli & Ginzburg, 2014). To compare coherence and trophic level we generated 1000 null model networks with the fitted parameters of the metaweb assembly model. Then we calculated the 99% confidence interval using the 0.5% and 99.5% quantiles of the distribution of *Q*. We also calculated the CI for the mean trophic level.

Another property related to stability is modularity, since the impacts of a perturbation are retained within modules minimizing impacts on the food web (Fortuna et al., 2010; Grilli, Rogers, & Allesina, 2016). It measures how strongly sub-groups of species interact between them compared with the strength of interaction with other sub-groups (Newman & Girvan, 2004). These sub-groups are called modules. To find the best partition, we used a stochastic algorithm based on simulated annealing (Reichardt & Bornholdt, 2006). Simulated annealing allows maximizing modularity without getting trapped in local maxima configurations (Roger Guimerà & Nunes Amaral, 2005). As the simulated annealing algorithm is stochastic we estimated modularity as the mean of 100 repetitions. To assess the significance of our networks we calculated the 99% confidence intervals based on 1000 null model networks as previously described.

Finally, we calculated the average of the maximal real part of the eigenvalues of the jacobian (Grilli, Rogers, & Allesina, 2016) for randomly parametrized systems, keeping fixed the predator-prey (sign) structure. This is a measure related to quasi sign-stability (QSS) that is the proportion of randomly parametrized systems that are locally stable (Allesina & Pascual, 2008). We sampled 1000 jacobians to estimate the maximal real part of the eigenvalues and withhold the average, we repeat this procedure for each of the 1000 null model networks and estimated the 99% confidence intervals as described earlier.

To show the results graphically we calculated the deviation for each metric, which correspond to the 99% confidence intervals for the metric’s value in the assembly null model. We define the mid-point

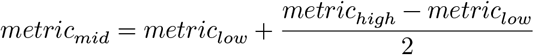

Then the deviation of the observed value of the real web is calculated

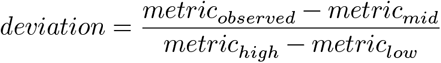

A deviation value outside of [-0.5, 0.5] indicates that the value is outside of the 99% confidence interval. See the Supporting Information for formulas and more details about these metrics.

### Motifs

We considered the abundance of sub-networks that deviates significantly from a null model network, which are called motifs (Milo et al., 2002). In practice, sub-networks are generally called motif without taking into account the mentioned condition. During the assembly process, motifs that are less dynamically stable tend to disappear from the food web (Borrelli, 2015; Borrelli et al., 2015). Furthermore, different motifs patterns could also be the result of habitat filtering (Dekel, Mangan, & Alon, 2005; Baldassano & Bassett, 2016).

We analyzed here the four three-species motifs that have been most studied theoretically and empirically in food webs (Prill, Iglesias, & Levchenko, 2005; Stouffer, Camacho, Jiang, & Nunes Amaral, 2007; Baiser, Elhesha, & Kahveci, 2016) (Figure 2). The four three-species motifs are: apparent competition, where two preys share a predator; exploitative competition, where two predators consume the same prey; omnivory, where predators feed at different trophic levels; and tri-trophic chain, where the top predator consumes an intermediate predator that consumes a basal prey (Figure 2). These are the most common motifs present in food webs (Borrelli, 2015; Monteiro & Del Bianco Faria, 2017). We compared the frequency of these motifs to 1000 null model networks using the 99% confidence intervals, and deviation as previously described.

**Figure 2:**
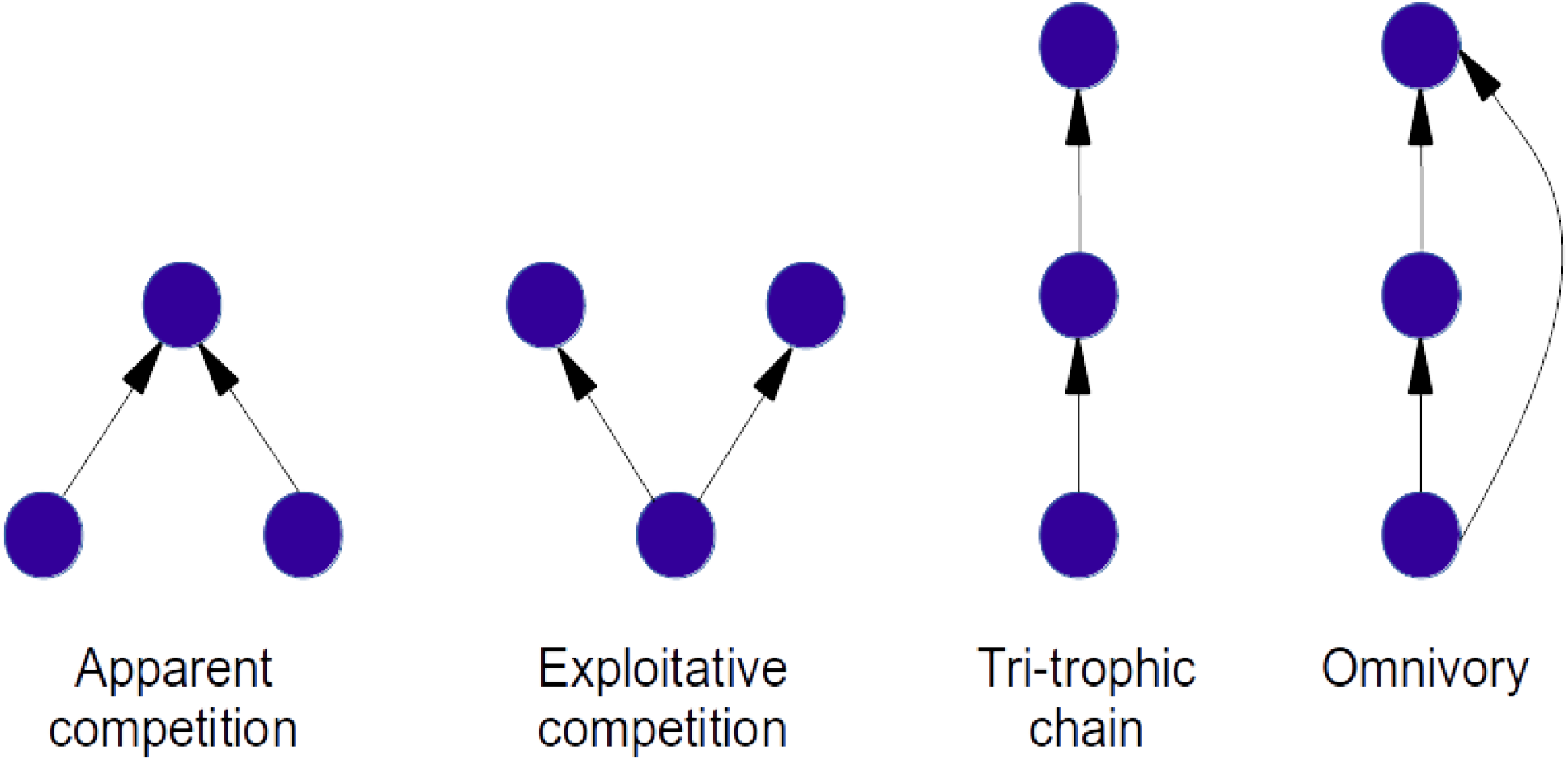
The four three-species motifs analysed: apparent competition, exploitative competition, tri-trophic chain, and omnivory. Motifs are three-node sub-networks. These four Motifs have been explored both theoretically and empirically in ecological networks and are the most common found in food webs

### Topological roles

To detect the process of habitat filtering or dispersal limitation in local food webs we calculated topological roles, which characterize how many trophic links are conducted within their module and/or between modules (Roger Guimerà & Nunes Amaral, 2005; Kortsch, Primicerio, Fossheim, Dolgov, & Aschan, 2015). Theoretical and empirical results suggest these roles are related to species traits, such as niche breadth, environmental tolerance, apex position in local communities and motility (Dupont & Olesen, 2009; Rezende, Albert, Fortuna, & Bascompte, 2009; R. Guimerà et al., 2010; Borthagaray, Arim, & Marquet, 2014; Kortsch, Primicerio, Fossheim, Dolgov, & Aschan, 2015).

We determined topological roles using the method of functional cartography (Roger Guimerà & Nunes Amaral, 2005), which is based on module membership (See Supporting Information for more details). There are four roles: *Hub connectors* that have a high number of between module links; *Module connectors* have a low number of links mostly between modules; *Module hubs* have a high number of links inside its module; *Module specialists* have a low number of links inside its module.

We estimated the roles for empirical networks and for 20 realizations of each assembly model network. To test if the proportion of species’ roles changed between the empirical and each of the realizations of the model we performed a Pearson’s Chi-squared test with simulated p-value based on 10000 Monte Carlo replicates.

All analyses and simulations were performed in R version 4.1.1 (R Core Team, 2017), using the igraph package version 1.2.6 (Csardi & Nepusz, 2006) for motifs, the package multiweb for topological roles, *Q* and other network metrics (Saravia, 2019), and the package meweasmo for the metaweb assembly model (Saravia, 2020).

## Results

A general description of all networks using the structural properties including metawebs is presented in appendix table S3. For the Antarctic metaweb, the differences in the number of species (size) between local food webs and the metaweb are greater than for the other metawebs. The metawebs of Florida Islands and Adirondacks’ Lakes have similar sizes and both are smaller and have higher connectance than the Antarctic metaweb. Thus there is a wide range of local food web sizes (13 to 435), number of links (17 to 1978), and connectance (0.01 to 0.29) in our dataset.

We found almost no differences between the assembly null model and the local food webs for trophic coherence (Q) except for E1 Island, which exhibited a lower value, hence, more stable (Figure 3, Table S5). The mean maximal eingenvalue (MEing) was also not different except for Weddell Sea, which has a lower value resulting in an increased local stability, and four local webs from the Lakes dataset (Briddge Brook Lake, Chub Pond, Hoel Lake, Long Lake) which have a higher MEing and lower stability than the model (Figure 3, Table S5). Only Weddell Sea and E1 Island were significantly different for mean trophic level (TL) (Figure 4, Table S4), only E1 Island have a lower TL, this should be the expected pattern if dynamical stability constraints where acting. For modularity we found only two local food webs different, Chub Pond, from Lakes metaweb, that is less modular than the model and Island E9 which is more modular (Figure 4, Table S5).

**Figure 3:**
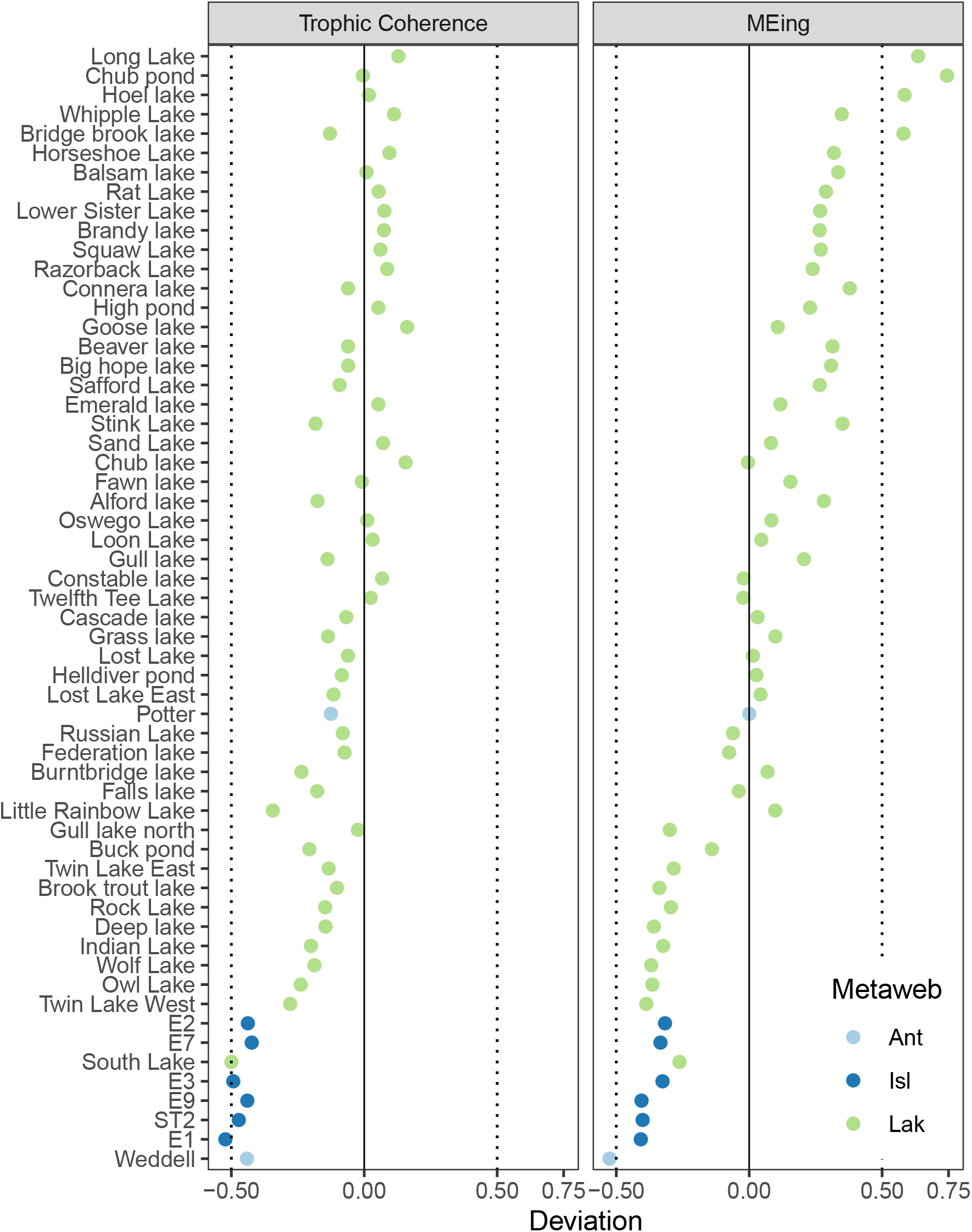
Trophic coherence (Q) and mean of maximal eingenvalue (MEIng) comparison for local empirical networks (dots) and assembly null model networks. We ran 1000 simulations of the metaweb assembly model fitted to local networks to build the 99% confidence intervals of the metric and calculated the deviation; a value outside -0.5,0.5 interval (vertical dotted lines) indicates that the value is outside of the 99% confidence interval. Colors represent metawebs to which local food webs belong, where *Ant* is the Antarctic, *Isl* is the Islands, and *Lak* the lakes metaweb.

**Figure 4:**
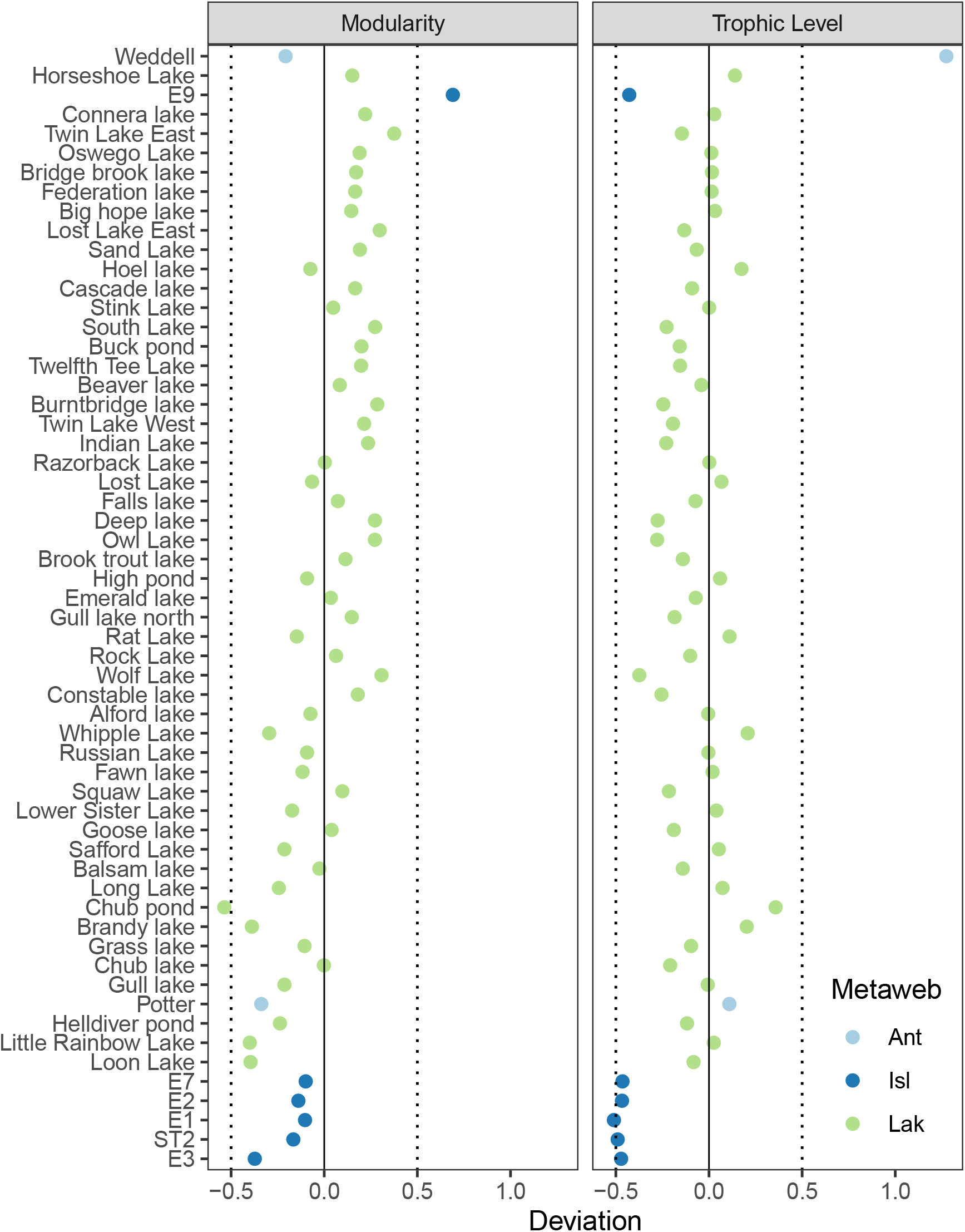
Modularity and mean trophic level comparison for local empirical networks (dots) and assembly null model networks. We ran 1000 simulations of the metaweb assembly model fitted to local networks to build the 99% confidence intervals of the metric and calculated the deviation; a value outside -0.5,0.5 interval (vertical dotted lines) indicates that the value is outside of the 99% confidence interval. Colours represent metawebs to which local food webs belong, where *Ant* is the Antarctic, *Isl* is the Islands, and *Lak* the lakes metaweb.

Comparing the motifs generated from the metaweb assembly null model, 9 of 58 (16%) networks showed at least one significant motif over-representation and only one (Weddell Sea) showed motifs under-representation (Figure 5, Table S6). The Hoel lake network was the only one that showed over-representation for all motifs. Long lake showed only omnivory over-representation, and 5 more have only 1 motif (not omnivory) over-representation. Apparent competition and exploitative competition were the most over-represented motifs (6 and 5 times).

**Figure 5:**
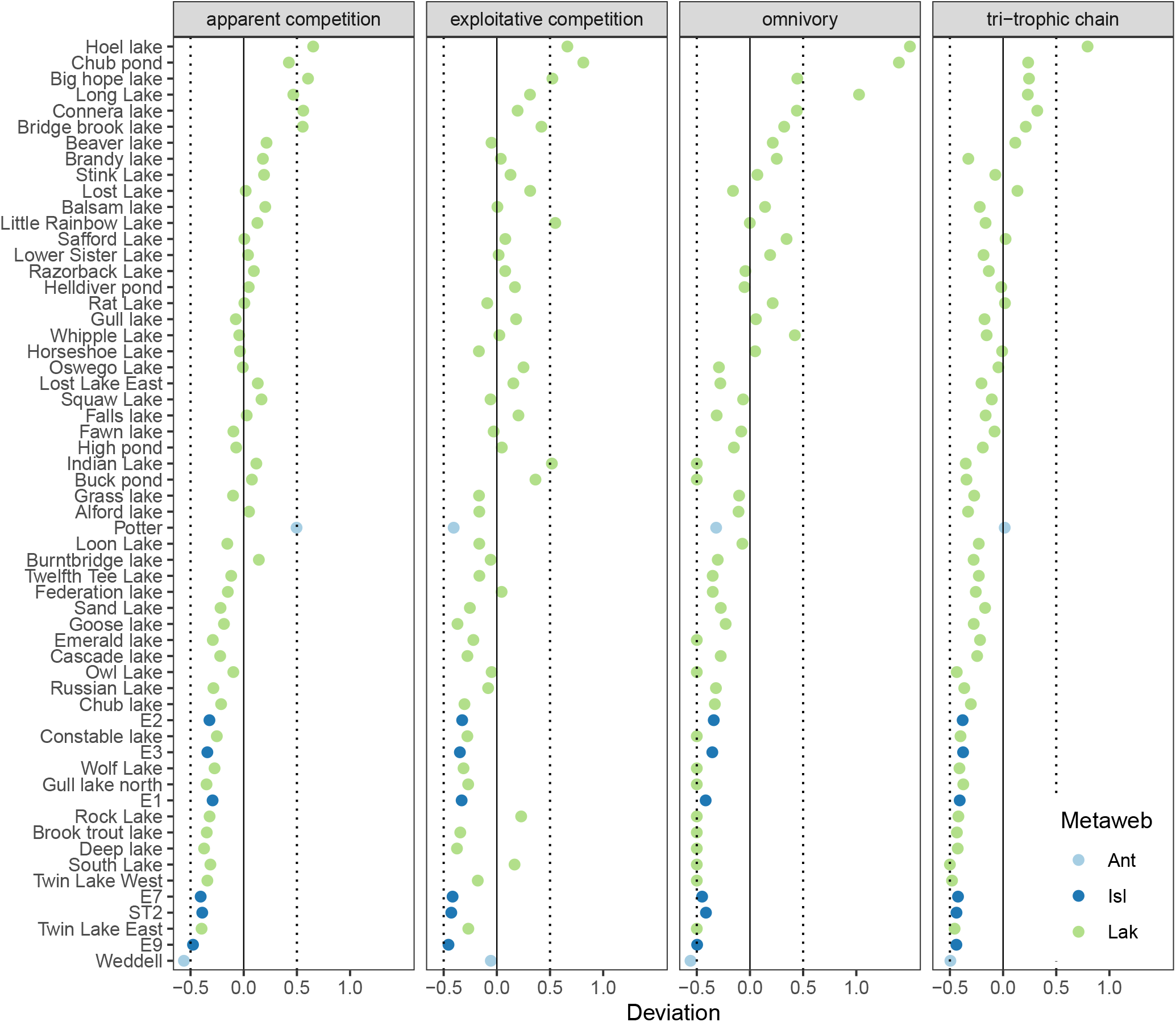
Motifs’ abundance comparison for local empirical networks (dots) and assembly null model networks. We ran 1000 simulations of the metaweb assembly model fitted to local networks to build the 99% confidence intervals of the metric and calculated the deviation; a value outside -0.5,0.5 interval (vertical dotted lines) indicates that the value is outside of the 99% confidence interval. Colors represent metawebs to which local food webs belong, where *Ant* is the Antarctic, *Isl* is the Islands, and *Lak* the lakes metaweb.

The proportions of topological roles were similar to the metaweb assembly model; across the 20 realizations of the assembly model, between 3 and 10 out of 58 local (5-17%) were different at 1% significant level (Table S8). Figure 6 shows the proportions for the Antarctic and Islands metawebs for one realization of the model and Figure S5 shows the proportions for the Lakes metaweb; we added the topological role proportions for the corresponding metaweb in each case to visually compare with both the empirical and model food webs. The only food web that showed consistent differences with the model was Potter Cove (100% of the realizations), the second the Island E9 with 60% and the third the Weddell Sea food web that showed differences 50% of the time (Table S9).

**Figure 6:**
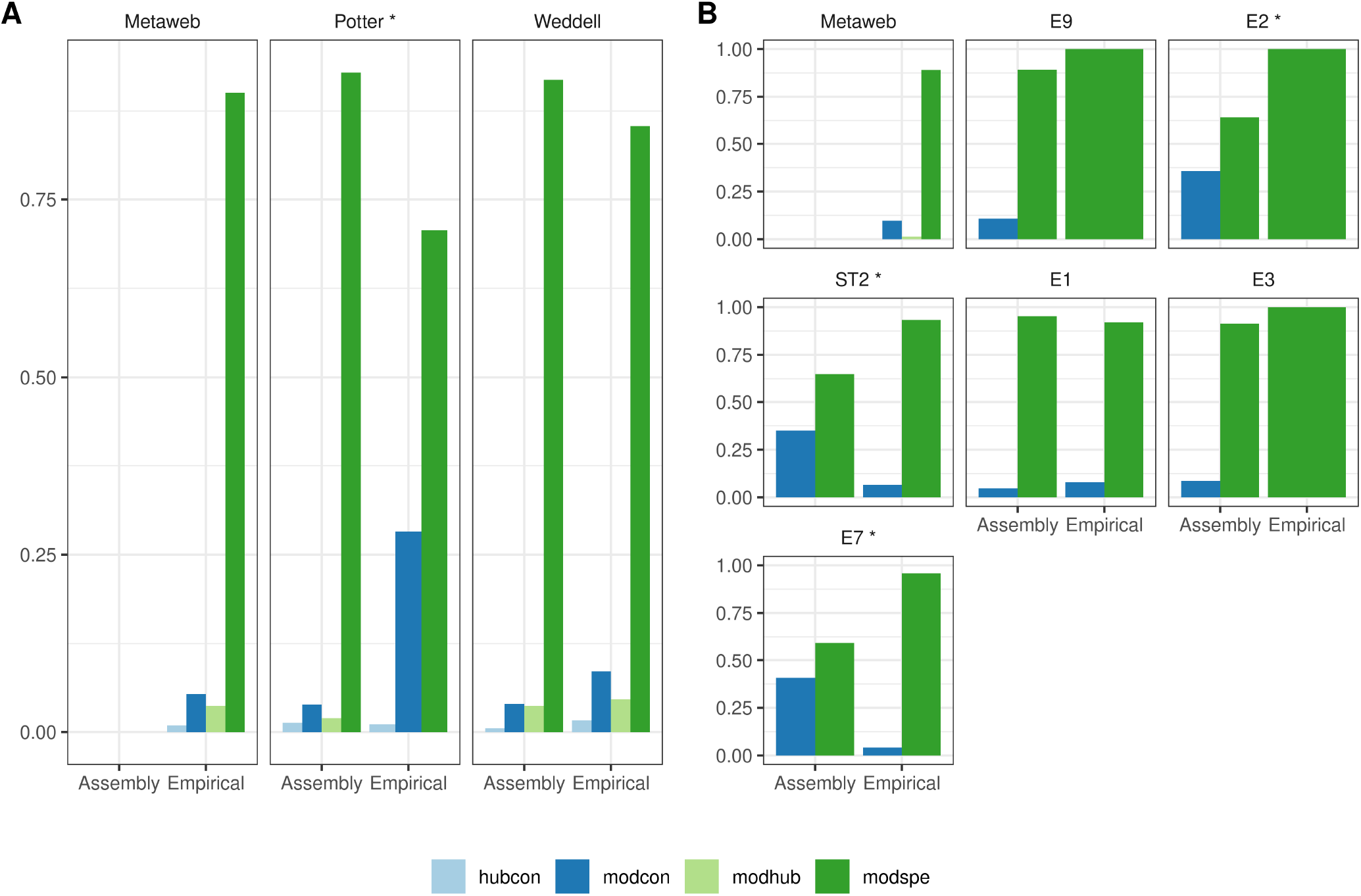
Topological roles proportions for local empirical networks and metawebs compared with assembly null model for the Antarctic (A) and Islands metawebs (B). The topological roles are: *Hub connectors* have a high number of between module links; *Module connectors* have a low number of links mostly between modules; *Module hubs* have a high number of links inside its module; *Module specialists* have a low number of links inside its module. Plots marked with ‘*’ are different from the null model at 1% level

## Discussion

We hypothesised that, if local processes like dynamical stability determine food web structure, then we should observe a consistent difference in network properties between real webs and model webs randomly assembled from the regional metaweb. Contrary to our expectations, we found that most structural properties did not differ significantly between real and randomly assembled webs. We investigated network properties associated with dynamical stability (trophic coherence, modularity, MEing, motifs). Although we found differences for some local food webs, there was not a general pattern, and real web properties did not display a consistent tendency to deviate in the direction predicted by theory. We also investigated topological roles, which we expected to change due to habitat filtering and dispersal limitation. However, with the exception of the two Antarctic webs, we found a similar lack of difference between real and model webs. These results suggest that — for the metrics we considered — food webs are mainly shaped by metaweb structure, and we did not find good evidence for the influence of local dynamics.

Local food webs are expected to have relatively few trophic levels (Richard J. Williams, Berlow, Dunne, Barabási, & Martinez, 2002; Borrelli & Ginzburg, 2014). Different hypotheses have been posed to explain this pattern: the low efficiency of energy transfer between trophic levels, predator size, predator behaviour, and consumer diversity (Young et al., 2013). Recently, it has been proposed that maximum trophic level could be related to productivity and ecosystem size depending on the context but related to energy fluxes that promote omnivory (Ward & McCann, 2017). Our results of mostly no differences with the randomly assembled webs, do not invalidate these previous hypotheses but point out that the mechanisms may not be acting at the scale of the assembly process.

We expected modularity to differ between real and randomly assembled webs both due to the influence of habitat heterogeneity (Krause, Frank, Mason, Ulanowicz, & Taylor, 2003; Rezende, Albert, Fortuna, & Bascompte, 2009) and modularity’s stability-enhancing effects. Recent studies suggest that modularity increases local stability, and this effect is stronger the more complex the network is (Stouffer & Bascompte, 2011). This suggests that modularity should be higher in real webs than random webs. However, the effect on stability also strongly depends on the interaction strength configuration (Grilli, Rogers, & Allesina, 2016) and the existence of external perturbations (Gilarranz, Rayfield, Liñán-Cembrano, Bascompte, & Gonzalez, 2017). We found in most cases no significant difference in modularity, which means that the species participating in the modules could change due to differences in habitats, but the strength of the modules in terms of number of within and between interactions is the same as the observed in the assembly model.

Due to dynamical stability constraints, we expected real webs to have a lower maximum eigenvalue (*MEing*) and higher trophic coherence (*Q*) than randomly assembled webs. However, only the Weddell Sea followed this expectation (i.e. greater stability, lower *MEing*), and four local food webs belonging to the Lakes metaweb showed the opposite pattern (i.e. lower stability, higher *MEing*). Thus, although this evidence is not conclusive concerning the importance of dynamical stability in the assembly of food webs, the structure of the local food webs examined here seems to be a consequence of the metaweb structure.

We also expected real webs to have a higher frequency of stability-enhancing motifs than randomly assembled webs. Specifically, we expected an over-representation of tri-trophic chains, exploitative competition, and apparent competition (Borrelli, 2015). Some Lakes food webs had an an over-representation of stability-enhancing motifs, but there was not a consistent pattern. Further, Lakes webs with an over-representation of stability-enhancing motifs often had low local stability (measured as high MEing). The omnivory motif can either enhance or diminish stability, depending on the context (Monteiro & Faria, 2016). In our study, omnivory seemed to be destabilizing: food webs with an over-representation of it (Chub Pond, Hoel Lake and Long Lake) were also those with significant lower local stability (higher MEing); and Weddell Sea, which had the most significant higher local stability result, also had the most significant under-representation of omnivory. Food webs are more than the sum of their three-species modules (Cohen, Schittler, Raffaelli, & Reuman, 2009), which is exemplified by the contradictory results for the Weddell Sea food web: high mean trophic level, enhancing stability, but an under-representation of omnivory and apparent competition motifs.

The relative proportions of topological roles were similar between real and randomly assembled webs except in two Antarctic food webs (Weddell Sea and Potter Cover) and some from the Islands metaweb (E2, E5, E9). These differences could reflect real differences between the habitats of the local webs and the metaweb. For example, in the metaweb, the Antarctic cod (*Notothenia coriiceps*) is a module hub (a species with most of its links within its module), but in Potter Cove, it is a super-generalist. In the Antarctic web, similar to observations in other Arctic and Caribbean marine food webs (Rezende, Albert, Fortuna, & Bascompte, 2009; Kortsch, Primicerio, Fossheim, Dolgov, & Aschan, 2015), modules typically correspond to a particular habitat, e.g., a benthic module, a pelagic module, etc. Consequently, given that local webs are smaller than the metaweb and could cover a particular habitat, habitat filtering (a local process that is not included in the random assembly model) could plays a large role. In contrast, the Lakes local food webs seem to be covered by similar local habitats.

The lack of difference between real and randomly assembled webs raises the possibility that, for these commonly used metrics that we measured, the values they take in local food webs merely reflect the properties of the metaweb. That does not mean that they are indeed the result of random assembly, nor that local processes like dynamical constraints do not act in reality. Rather, it suggests that, without further evidence, one cannot take for granted that these metrics reflect dynamical constraints (see Zhang (2020) for a similar argument regarding species co-occurrence patterns). This is particularly true given that the metrics associated with dynamical constraints can also be subordinate to other more fundamental structure-determining forces. For example, high trophic coherence means that the trophic levels are fairly distinct and there is little to no omnivory. Trophic coherence is associated with high stability; however, it has also been been modelled as a consequence of the relative strength of competition versus width of consumption niche, all mediated by a physiological trait such as body size (Loeuille & Loreau, 2005). Further, adaptive foraging (Heckmann, Drossel, Brose, & Guill, 2012) also leads to the emergence of trophic coherence in theoretical assembly models (Drossel, Higgs, & Mckane, 2001), which has the side-effect of lowering connectance and hence increasing stability (Beckerman, Petchey, & Warren, 2006).

Assuming that population dynamics does indeed play some role in determining food web structure, why then did we not observe its signal? One possible reason is that the effect of dynamics on the local network properties also manifests on regional scales. The metaweb structure is an aggregation of local webs (Ricklefs, 1987; Araújo & Rozenfeld, 2014); therefore, if dynamical constraints prevent certain network structures from occurring in local webs, then they are also prevented in the metaweb. However, while it is obviously true that local processes influence metaweb structure, we still expected to see differences between real webs and those randomly assembled from the metaweb. Local food webs differ from one another due to e.g., historical contingencies, like the stochastically determined order of arrival of species. These differences between local webs provide the species and structural diversity in the metaweb upon which non-Darwinian selection process is theorised to act (Borrelli, 2015). It is important to note that a species that is stabilising in one food web can be destabilising in another. In contrast, the metaweb describes species co-occurrences and interaction possibilities that do not occur in reality, including those that are presumably precluded due to dynamical constraints. Therefore, if the non-Darwinian selection process is true, we should expect to see differences between real and randomly assembled webs, not only due to these historical contingencies, but also due to subsequent dynamical constraints that precluded certain co-occurrences and interactions from occurring.

Another possible reason why we did not observe the expected difference between real and randomly assembled webs is that these commonly used metrics are too coarse to detect the signal of dynamical constraint. We chose popular structural metrics that have been associated with stability in the literature; however, stability also depends on interaction strengths. For example, generalisations about the relationship between modularity and stability cannot be made without first characterising the distribution of interaction strengths (Grilli, Rogers, & Allesina, 2016), which were unknown for our webs. Further, given that most predator-prey interactions are weak (K. McCann, Hastings, & Huxel, 1998; Neutel, Heesterbeek, & Ruiter, 2002), the structure of the species-rich food webs we investigated might mask the importance of the few strong links. Theoretical predictions relating stability to structure also depend on the particulars of the community and the type of perturbation considered (Cenci, Song, & Saavedra, 2018). It is known that small changes in network structure can have large effects on food web stability (Fox, 2006). It is also known that the sum of effects of positive and negative feedback loops that determine stability can interact in counterintuitive ways (Hosack, Li, & Rossignol, 2009). Therefore, it may not be possible to reduce those complex interactions into simple structural metrics.

In conclusion, we found that the commonly used metrics of network structure do not differ between real food webs and model webs randomly assembled from the regional metaweb. This suggests that evolutionary and metacommunity assembly processes are more important to these aspects of food web structure than local dynamics. However, this kind of analysis needs to be expanded to other regions and habitat types to confirm whether or not this is a general pattern.

## Acknowledgements

We are grateful to the National University of General Sarmiento for financial support (Project 30/1139). LAS would like to thank Susanne Kortsch that shared with us her source code for topological analysis and figures. This work was partially supported by a grant from CONICET (PIO 144-20140100035-CO).

## Authors’ contributions

LAS, TIM, MDT and FRM conceived the ideas and designed methodology; TIM and LAS collected the data; LAS wrote the code; LAS, TIM and NPK analysed the data; NPK, LAS and TIM led the writing of the manuscript. All authors contributed critically to the drafts and gave final approval for publication.

## Data Availability Statement

The source code and data is available at zenodo https://doi.org/10.5281/zenodo.3370022 and Github https://github.com/lsaravia/MetawebsAssembly/.

