## Supplemental Information for "Ecological network assembly: how the regional metaweb influences local food webs"

### Supplementary Information

#### Formulas for the network properties

To describe food webs as networks each species is represented as a node or vertex and the trophic interactions are represented as edges or links between nodes. These links are directed, from the prey to the predator, as the flow of energy and matter. Two nodes are neighbours if they are connected by an edge and the degree $k_i$ of node $i$ is the number of neighbours it has. The food web can be represented by an adjacency matrix $A = (a_{ij})$ where $a_{ij} = 1$ if species $j$ predates on species $i$, else is 0. Then $k_i^{in} = \sum_j a_{ji}$ is the number of preys of species $i$ or its in-degree, and $k_i^{out} = \sum_j a_{ij}$ is the number of predators of $i$ or its out-degree. The total number of edges is $L = \sum_{ij} a_{ij}$.

##### Trophic coherence

We first needed to estimate the trophic level of a node $i$, defined as the average trophic level of its preys plus 1. That is:

$$tp_i = 1 + \frac{1}{k_i^{in}} \sum_j a_{ij} tp_j \tag{1}$$

where $k_i^{in} = \sum_j a_{ji}$ is the number of preys of species $i$, basal species that do not have preys (then $k_i^{in} = 0$) are assigned a $tp = 1$. Then the trophic difference associated to each edge is defined as $x_{ij} = tp_i - tp_j$. The distribution of trophic differences, $p(x)$, has a mean $L^{-1} \sum_{ij} a_{ij} x_{ij} = 1$ by definition. Then the trophic coherence is measured by:

$$Q = \sqrt{\frac{1}{L} \sum_{ij} a_{ij} x_{ij}^2 - 1} \tag{2}$$

that is the standard deviation of the distribution of all trophic distances.

##### Modularity

The index of modularity was defined as:

$$M = \sum_s \left( \frac{I_s}{L} - \left( \frac{d_s}{2L} \right)^2 \right) \tag{3}$$

where $s$ is the number of modules, $I_s$ is the number of links between species in the module $s$, $d_s$ is the sum of degrees for all species in module $s$ and $L$ is the total number of links for the network.

##### Average of the maximal real part of the eigenvalues of the jacobian (MEing)

The Jacobian $J$, so-called community matrix (May, 1974), represents the population-level effect of a change in one species' density on any other species, including the dependence on its own density (self-regulation), at an equilibrium. A system is locally stable if the Jacobian $J$ has all its eigenvalues negative, thus the maximal eigenvalue has to be less than zero for a system to be locally stable. The signs of the elements of $J$ are given by the predator-prey structure of the food web, but the magnitude of the elements are unknown. Following previous analysis (Borrelli, 2015; Monteiro & Faria, 2016), we estimated the unknown magnitudes by drawing the predator-prey interactions from a uniform distribution ranging from -10 to 0, the prey-predator interactions from 0 to 0.1, and from 0 to -1 for the self-regulation effect. This implies that the predator effect on the prey is bigger than the effect of the prey on the predator, and that the self-regulation or self-damping effect, that scales the dynamic's return time, is generally smaller than the predator-prey effect. Besides other parametrizations are possible they give very similar results (not shown).

##### Topological roles

The roles are characterized by two parameters: the standardized within-module degree $dz$ and the among-module connectivity participation coefficient $PC$. The within-module degree is a z-score that measures how well a species is connected to other species within its module:

$$dz_i = \frac{k_{is} - \bar{k_s}}{\sigma_{ks}}$$

where $k_{is}$ is the number of links of species $i$ within its own module $s$, $\bar{k_s}$ and $\sigma_{ks}$ are the average and standard deviation of $k_{is}$ over all species in $s$. The participation coefficient $PC$ estimates the distribution of the links of species $i$ among modules; thus it can be defined as:

$$PC_i = 1 - \sum_s \frac{k_{is}}{k_i}$$

where $k_i$ is the degree of species $i$ (i.e. the number of links), $k_{is}$ is the number of links of species $i$ to species in module $s$. Due to the stochastic nature of the module detection algorithm, we made repeated runs of the algorithm (starting from 100 runs) until there were no statistical differences between the distributions of $PC_i$ and $dz_i$ in successive repetitions; to test such statistical difference we used the k-sample Anderson-Darling test (Scholz & Stephens, 1987). Then we calculated the mean and 95% confidence interval of $dz$ and $PC$.

To determine each species' role the $dz - PC$ parameter space was divided into four areas, modified from Guimerà & Nunes Amaral (2005), using the same scheme as Kortsch, Primicerio, Fossheim, Dolgov, & Aschan (2015). Two thresholds were used to define the species' roles: $PC = 0.625$ and $dz = 2.5$. If a species had at least 60% of links within its module then $PC < 0.625$, and if it also had $dz \geq 2.5$, thus it was classified as a module hub. This parameter space defines species with a relatively high number of links, the majority within its module. If a species had $PC < 0.625$ and $dz < 2.5$, then it was called a peripheral or specialist; this refers to a species with relatively few links, mostly within its module. Species that had $PC \geq 0.625$ and $dz < 2.5$ were considered module connectors, since they have relatively few links, mostly between modules. Finally, if a species had $PC \geq 0.625$ and $dz \geq 2.5$, then it was classified as a super-generalist or hub-connector because it has high between- and within-module connectivity.

#### Meta-web assembly model, steady state and additional checks

We performed exploratory simulations to determine if the last 100 time steps of 1000 simulated were enough to avoid the transient period and sample the attractor of the model. Figures S1 and S2 show that there are no differences in the dynamics after 1000 or 3000 time steps and that the transient is well over after 1000 time steps. We repeated these exploratory simulations with the three metawebs obtaining similar results.

Table S1: Range of parameters used in latin hypercubic sampling to simulate the metaweb assembly model. The range of each parameter is divided into into $n = 150000$ equal partitions and random data points are chosen such that each partition in each dimension is sampled once, i.e., ensuring coverage of the entire range.

| Parameter | Low | High |
|---|---|---|
| $c$ | 0.010 | 0.3 |
| $e$ | 0.001 | 0.5 |
| $se$ | 0.100 | 0.9 |

After fitting the model as explained in the main text we performed an additional validation of the model by doing simulations with the fitted parameters and checked that the number of species $S_e$ and connectance $C_e$ were inside the 99% confidence interval (CI) generated with 1000 simulations of the model (Table S4, Figures S3 & S4). Only in two cases (Bridge_brook_lake, Lak_Chub_pond) the connectance was higher than the CI.

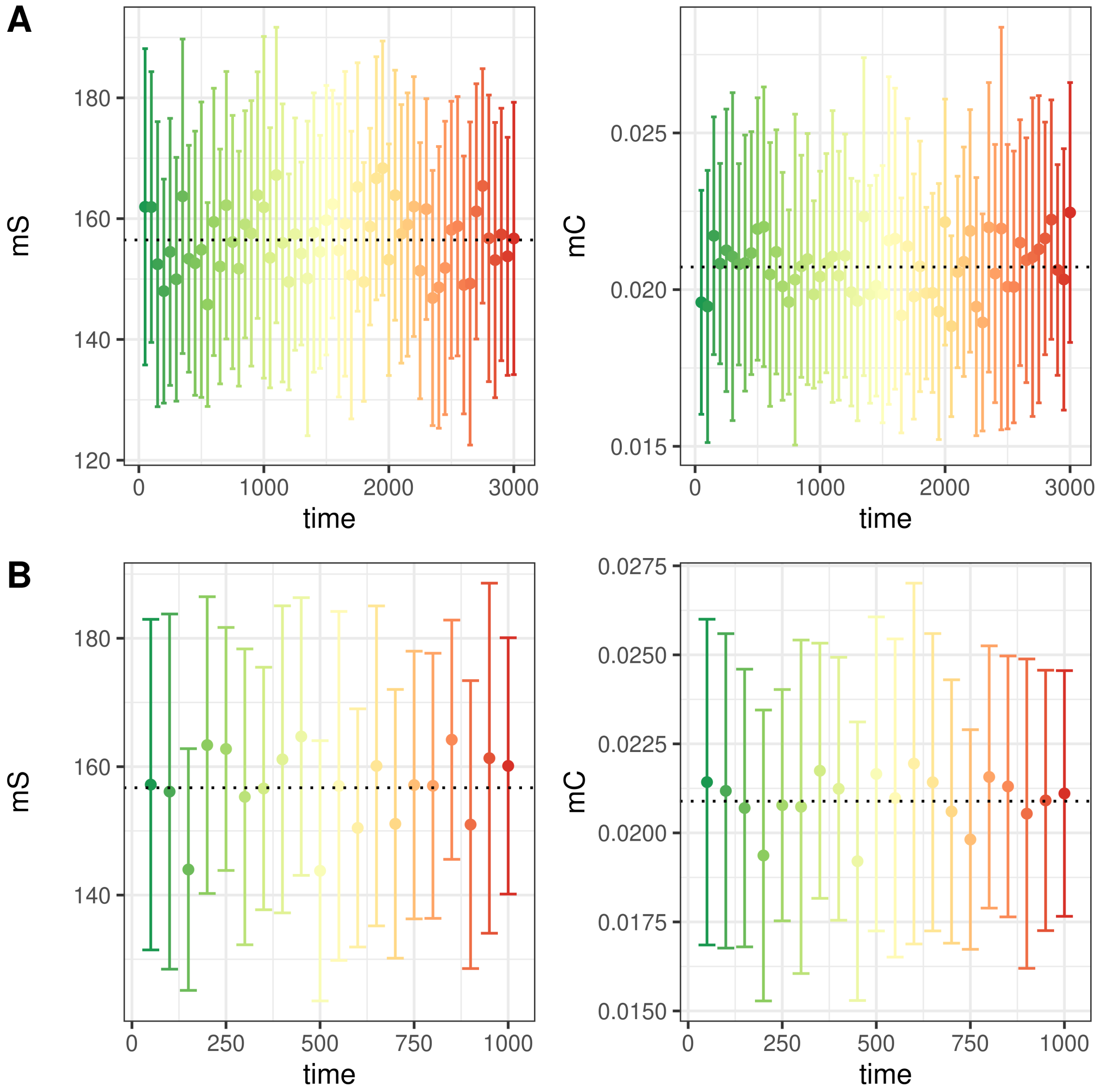


Figure S1: Windowed averages of a simulation of the Meta-web assembly model with the Antarctic metaweb and parameters near the fitted to Potter Cove food web, where $mS$ and $mC$ are windowed averages of the number of species and connectance, error bars are the standard deviation, and the dashed is line the overall mean using a window of 50 time steps. The parameters were $c$=0.02, $e$=0.05, $se$=0.70.

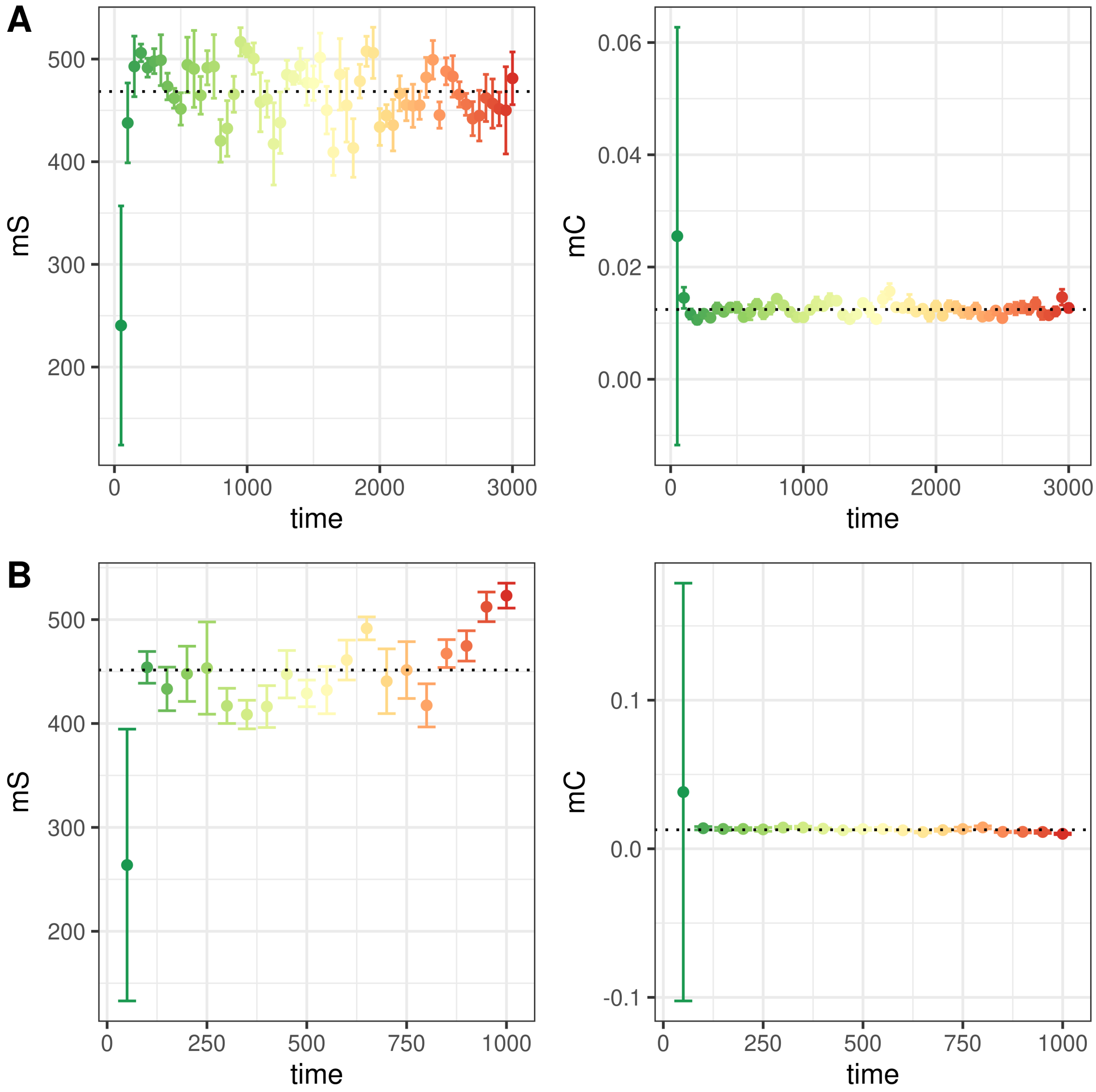


Figure S2: Windowed averages of a simulation of the Meta-web assembly model with the Antarctic metaweb and parameters near the fitted to Weddell Sea food web. Where $mS$ and $mC$ are windowed averages of the number of species and connectance, error bars are the standard deviation, and the dashed is line the overall mean using a window of 50 time steps. The parameters were $c$=0.03, $e$=0.016, $se$=0.10.

Table S2: Metaweb assembly model best fit of the local food webs for the 3 metawebs: Ant=Antarctic, Isl=Islands, Lak=Lakes; where $c$ is the colonization parameter, $e$ the extinction rate and $se$ the secondary extinction rate. S is the number of species, L the number of links, and C the connectance averaged over the last 100 time steps. $\alpha = c/e$ is the hypothesized distance to main land, and the *distance* the relative distance to the empirical values.

| Metaweb | Network | $c$ | $e$ | $se$ | S | L | C | $\alpha$ | distance |
|---|---|---|---|---|---|---|---|---|---|
| Ant | Potter | 0.0188 | 0.0552 | 0.7648 | 93.5050 | 346.0792 | 0.0396 | 0.3412 | 0.0349 |
| Ant | Weddell | 0.0271 | 0.0164 | 0.1016 | 450.3663 | 2312.5446 | 0.0114 | 1.6509 | 0.0973 |
| Isl | E1 | 0.0102 | 0.0319 | 0.4055 | 25.2673 | 116.2970 | 0.1822 | 0.3209 | 0.0108 |
| Isl | E2 | 0.0189 | 0.0397 | 0.5517 | 28.3267 | 161.1089 | 0.2008 | 0.4752 | 0.3226 |
| Isl | E3 | 0.0259 | 0.0764 | 0.6145 | 23.7030 | 115.9307 | 0.2063 | 0.3389 | 0.1973 |
| Isl | E7 | 0.0259 | 0.0764 | 0.6145 | 23.7030 | 115.9307 | 0.2063 | 0.3389 | 0.0143 |
| Isl | E9 | 0.0210 | 0.0293 | 0.4132 | 33.0099 | 205.0693 | 0.1882 | 0.7167 | 0.0851 |
| Isl | ST2 | 0.0189 | 0.0397 | 0.5517 | 28.3267 | 161.1089 | 0.2008 | 0.4752 | 0.1426 |
| Lak | Alford_lake | 0.0375 | 0.0735 | 0.7522 | 56.5545 | 224.9406 | 0.0703 | 0.5095 | 0.0102 |
| Lak | Balsam_lake | 0.0239 | 0.0503 | 0.3548 | 49.7426 | 196.7723 | 0.0795 | 0.4749 | 0.2383 |
| Lak | Beaver_lake | 0.0163 | 0.0363 | 0.4049 | 52.3366 | 210.6634 | 0.0769 | 0.4492 | 0.1167 |
| Lak | Big_hope_lake | 0.0160 | 0.0390 | 0.5759 | 56.1287 | 236.4752 | 0.0751 | 0.4094 | 0.1686 |
| Lak | Brandy_lake | 0.0122 | 0.0438 | 0.2648 | 32.5347 | 106.0396 | 0.1002 | 0.2794 | 0.2685 |
| Lak | Bridge_brook_lake | 0.0221 | 0.0304 | 0.2525 | 76.6040 | 388.0594 | 0.0661 | 0.7284 | 0.3280 |
| Lak | Brook_trout_lake | 0.0383 | 0.2806 | 0.1126 | 15.2772 | 19.6337 | 0.0841 | 0.1366 | 0.0189 |
| Lak | Buck_pond | 0.0357 | 0.1045 | 0.6141 | 39.3861 | 140.1980 | 0.0904 | 0.3417 | 0.0400 |
| Lak | Burntbridge_lake | 0.0317 | 0.0794 | 0.6234 | 52.9901 | 183.0495 | 0.0652 | 0.3991 | 0.0007 |
| Lak | Cascade_lake | 0.0288 | 0.0857 | 0.7663 | 35.0594 | 111.2871 | 0.0905 | 0.3357 | 0.0601 |
| Lak | Chub_lake | 0.0797 | 0.2744 | 0.5189 | 36.0099 | 83.0990 | 0.0641 | 0.2906 | 0.0007 |
| Lak | Chub_pond | 0.0239 | 0.0503 | 0.3548 | 49.7426 | 196.7723 | 0.0795 | 0.4749 | 0.4495 |
| Lak | Connera_lake | 0.0152 | 0.0280 | 0.6940 | 64.7723 | 290.7030 | 0.0693 | 0.5447 | 0.2997 |
| Lak | Constable_lake | 0.0505 | 0.1934 | 0.6342 | 31.8317 | 60.6040 | 0.0598 | 0.2611 | 0.0066 |
| Lak | Deep_lake | 0.0257 | 0.1429 | 0.7522 | 19.0198 | 28.2376 | 0.0781 | 0.1800 | 0.0065 |
| Lak | Emerald_lake | 0.0109 | 0.0562 | 0.8963 | 21.2970 | 50.6535 | 0.1117 | 0.1946 | 0.0752 |
| Lak | Falls_lake | 0.0357 | 0.1045 | 0.6141 | 39.3861 | 140.1980 | 0.0904 | 0.3417 | 0.0962 |
| Lak | Fawn_lake | 0.0122 | 0.0438 | 0.2648 | 32.5347 | 106.0396 | 0.1002 | 0.2794 | 0.1600 |
| Lak | Federation_lake | 0.0109 | 0.0562 | 0.8963 | 21.2970 | 50.6535 | 0.1117 | 0.1946 | 0.0608 |
| Lak | Goose_lake | 0.1494 | 0.4863 | 0.2929 | 39.9406 | 97.5050 | 0.0611 | 0.3071 | 0.0026 |
| Lak | Grass_lake | 0.0251 | 0.0612 | 0.6190 | 42.9901 | 155.2574 | 0.0840 | 0.4101 | 0.0271 |
| Lak | Gull_lake | 0.0251 | 0.0612 | 0.6190 | 42.9901 | 155.2574 | 0.0840 | 0.4101 | 0.2026 |
| Lak | Gull_lake_north | 0.0690 | 0.4868 | 0.8885 | 16.0198 | 25.0198 | 0.0975 | 0.1417 | 0.0021 |
| Lak | Helldiver_pond | 0.0357 | 0.1045 | 0.6141 | 39.3861 | 140.1980 | 0.0904 | 0.3417 | 0.1084 |
| Lak | High_pond | 0.0102 | 0.0495 | 0.3734 | 22.1188 | 51.9604 | 0.1062 | 0.2058 | 0.3070 |
| Lak | Hoel_lake | 0.0179 | 0.0323 | 0.3971 | 69.2970 | 321.3267 | 0.0669 | 0.5533 | 0.3943 |
| Lak | Horseshoe_Lake | 0.0239 | 0.0503 | 0.3548 | 49.7426 | 196.7723 | 0.0795 | 0.4749 | 0.2517 |
| Lak | Indian_Lake | 0.0380 | 0.1501 | 0.5492 | 34.2673 | 97.7327 | 0.0832 | 0.2535 | 0.0209 |
| Lak | Little_Rainbow_Lake | 0.0239 | 0.0503 | 0.3548 | 49.7426 | 196.7723 | 0.0795 | 0.4749 | 0.1365 |
| Lak | Long_Lake | 0.0152 | 0.0280 | 0.6940 | 64.7723 | 290.7030 | 0.0693 | 0.5447 | 0.2980 |
| Lak | Loon_Lake | 0.0288 | 0.0857 | 0.7663 | 35.0594 | 111.2871 | 0.0905 | 0.3357 | 0.0601 |
| Lak | Lost_Lake | 0.0122 | 0.0438 | 0.2648 | 32.5347 | 106.0396 | 0.1002 | 0.2794 | 0.3530 |
| Lak | Lost_Lake_East | 0.0151 | 0.0503 | 0.3573 | 40.0891 | 129.9703 | 0.0809 | 0.2991 | 0.0235 |
| Lak | Lower_Sister_Lake | 0.0288 | 0.0857 | 0.7663 | 35.0594 | 111.2871 | 0.0905 | 0.3357 | 0.2360 |
| Lak | Oswego_Lake | 0.0122 | 0.0438 | 0.2648 | 32.5347 | 106.0396 | 0.1002 | 0.2794 | 0.2099 |
| Lak | Owl_Lake | 0.0476 | 0.2041 | 0.5382 | 29.9901 | 70.9109 | 0.0788 | 0.2334 | 0.0007 |

| Metaweb | Network | $c$ | $e$ | $se$ | S | L | C | $\alpha$ | distance |
|---|---|---|---|---|---|---|---|---|---|
| Lak | Rat_Lake | 0.0239 | 0.0503 | 0.3548 | 49.7426 | 196.7723 | 0.0795 | 0.4749 | 0.2718 |
| Lak | Razorback_Lake | 0.0357 | 0.1045 | 0.6141 | 39.3861 | 140.1980 | 0.0904 | 0.3417 | 0.1258 |
| Lak | Rock_Lake | 0.0655 | 0.3465 | 0.7131 | 22.0099 | 42.9505 | 0.0887 | 0.1890 | 0.0021 |
| Lak | Russian_Lake | 0.0400 | 0.1631 | 0.6934 | 23.9703 | 58.0099 | 0.1010 | 0.2453 | 0.0467 |
| Lak | Safford_Lake | 0.0251 | 0.0612 | 0.6190 | 42.9901 | 155.2574 | 0.0840 | 0.4101 | 0.2781 |
| Lak | Sand_Lake | 0.0246 | 0.1060 | 0.4420 | 28.1881 | 77.7030 | 0.0978 | 0.2323 | 0.0282 |
| Lak | South_Lake | 0.0418 | 0.2388 | 0.6839 | 22.2574 | 36.7624 | 0.0742 | 0.1751 | 0.0119 |
| Lak | Squaw_Lake | 0.0357 | 0.1045 | 0.6141 | 39.3861 | 140.1980 | 0.0904 | 0.3417 | 0.1084 |
| Lak | Stink_Lake | 0.0239 | 0.0503 | 0.3548 | 49.7426 | 196.7723 | 0.0795 | 0.4749 | 0.2140 |
| Lak | Twelfth_Tee_Lake | 0.0496 | 0.2142 | 0.3354 | 31.0693 | 82.0297 | 0.0850 | 0.2317 | 0.0162 |
| Lak | Twin_Lake_East | 0.0163 | 0.1425 | 0.8147 | 12.9604 | 17.0198 | 0.1013 | 0.1145 | 0.0079 |
| Lak | Twin_Lake_West | 0.0253 | 0.0919 | 0.8725 | 26.0000 | 59.9307 | 0.0887 | 0.2749 | 0.0012 |
| Lak | Whipple_Lake | 0.0122 | 0.0438 | 0.2648 | 32.5347 | 106.0396 | 0.1002 | 0.2794 | 0.2463 |
| Lak | Wolf_Lake | 0.0154 | 0.0668 | 0.5350 | 26.7624 | 41.0495 | 0.0573 | 0.2299 | 0.0102 |

Table S3: Topological metrics of all empirical (local) food webs including metawebs, where **Ant** is the Antarctic, **Lak** the Lakes, **Isl** the islands metaweb. Size is the number of species $S$, Links is the number of trophic links $L$, Connectance $C$. Trophic coherence $Q$, mean trophic level $TL$, Modularity $MO$, and mean maximal eingenvalue $MEing$ are network metrics explained in the main text.

| Metaweb | Network | Size | Links | Connectance | Q | TL | MO | MEing |
|---|---|---|---|---|---|---|---|---|
| Ant | Potter | 92 | 325 | 0.0384 | 0.5230 | 2.1414 | 0.3601 | 2.1620 |
| Ant | Weddell | 435 | 1978 | 0.0105 | 0.5829 | 2.7908 | 0.4586 | 3.6264 |
| Ant | Meta | 846 | 6897 | 0.0096 | 0.7696 | 2.2423 | 0.4786 | 6.7254 |
| Isl | E2 | 31 | 280 | 0.2914 | 0.7507 | 2.6022 | 0.1601 | 4.0469 |
| Isl | E3 | 24 | 148 | 0.2569 | 0.6157 | 2.3143 | 0.1334 | 2.9558 |
| Isl | E7 | 24 | 118 | 0.2049 | 0.7047 | 2.2584 | 0.1765 | 2.8429 |
| Isl | E9 | 33 | 224 | 0.2057 | 0.7660 | 2.8061 | 0.2437 | 5.3367 |
| Isl | ST2 | 30 | 208 | 0.2311 | 0.7006 | 2.3627 | 0.1561 | 3.6717 |
| Isl | Meta | 155 | 5114 | 0.2129 | 0.8471 | 2.8653 | 0.1565 | 13.5643 |
| Lak | Twin_Lake_East | 13 | 17 | 0.1006 | 0.3333 | 1.4872 | 0.3962 | -0.1111 |
| Lak | Brook_trout_lake | 15 | 19 | 0.0844 | 0.3244 | 1.5111 | 0.2909 | -0.1133 |
| Lak | Gull_lake_north | 16 | 25 | 0.0977 | 0.4121 | 1.4732 | 0.2960 | -0.0735 |
| Lak | Deep_lake | 19 | 28 | 0.0776 | 0.3164 | 1.3977 | 0.3680 | -0.0682 |
| Lak | South_Lake | 22 | 36 | 0.0744 | 0.0000 | 1.4091 | 0.3492 | 0.0417 |
| Lak | Wolf_Lake | 27 | 42 | 0.0576 | 0.2583 | 1.2798 | 0.3759 | 0.0100 |
| Lak | Rock_Lake | 22 | 43 | 0.0888 | 0.2859 | 1.6201 | 0.2750 | 0.0069 |
| Lak | Federation_lake | 22 | 57 | 0.1178 | 0.3508 | 1.7339 | 0.3201 | 0.5067 |
| Lak | Emerald_lake | 22 | 58 | 0.1198 | 0.4828 | 1.6364 | 0.2519 | 0.9974 |
| Lak | Twin_Lake_West | 26 | 60 | 0.0888 | 0.1653 | 1.4846 | 0.3664 | 0.0968 |
| Lak | Constable_lake | 32 | 61 | 0.0596 | 0.4090 | 1.4107 | 0.3671 | 0.9281 |
| Lak | Russian_Lake | 24 | 61 | 0.1059 | 0.3269 | 1.6815 | 0.1983 | 0.8123 |
| Lak | Owl_Lake | 30 | 71 | 0.0789 | 0.2010 | 1.4212 | 0.3929 | 0.1351 |
| Lak | Sand_Lake | 29 | 82 | 0.0975 | 0.4567 | 1.6248 | 0.3412 | 1.0770 |
| Lak | Chub_lake | 36 | 83 | 0.0640 | 0.4514 | 1.4615 | 0.3048 | 1.1629 |
| Lak | Twelfth_Tee_Lake | 31 | 83 | 0.0864 | 0.4122 | 1.5079 | 0.3736 | 0.8816 |
| Lak | High_pond | 24 | 87 | 0.1510 | 0.4843 | 1.8835 | 0.1904 | 1.3283 |
| Lak | Goose_lake | 40 | 98 | 0.0612 | 0.4375 | 1.4658 | 0.3362 | 1.5594 |

| Metaweb | Network | Size | Links | Connectance | Q | TL | MO | MEing |
|---|---|---|---|---|---|---|---|---|
| Lak | Indian_Lake | 35 | 102 | 0.0833 | 0.2173 | 1.4590 | 0.3919 | 0.2044 |
| Lak | Cascade_lake | 35 | 118 | 0.0963 | 0.3983 | 1.6400 | 0.3800 | 1.3142 |
| Lak | Loon_Lake | 35 | 118 | 0.0963 | 0.4436 | 1.6612 | 0.1782 | 1.4555 |
| Lak | Brandy_lake | 30 | 121 | 0.1344 | 0.4179 | 1.9132 | 0.1216 | 1.7252 |
| Lak | Fawn_lake | 32 | 122 | 0.1191 | 0.3640 | 1.7077 | 0.2741 | 1.4517 |
| Lak | Whipple_Lake | 32 | 136 | 0.1328 | 0.4370 | 1.9118 | 0.2001 | 1.9576 |
| Lak | Lost_Lake_East | 41 | 137 | 0.0815 | 0.2734 | 1.5691 | 0.4095 | 1.2536 |
| Lak | Oswego_Lake | 33 | 138 | 0.1267 | 0.3659 | 1.6789 | 0.3235 | 1.2401 |
| Lak | Lost_Lake | 31 | 148 | 0.1540 | 0.3244 | 1.8110 | 0.2510 | 1.1592 |
| Lak | Falls_lake | 39 | 152 | 0.0999 | 0.2954 | 1.6161 | 0.3476 | 1.2022 |
| Lak | Buck_pond | 41 | 153 | 0.0910 | 0.1974 | 1.5301 | 0.3843 | 0.8899 |
| Lak | Lower_Sister_Lake | 37 | 161 | 0.1176 | 0.4090 | 1.7332 | 0.2442 | 1.9143 |
| Lak | Grass_lake | 44 | 165 | 0.0852 | 0.3979 | 1.6423 | 0.2957 | 1.6909 |
| Lak | Helldiver_pond | 41 | 169 | 0.1005 | 0.3787 | 1.5881 | 0.2446 | 1.3853 |
| Lak | Squaw_Lake | 41 | 169 | 0.1005 | 0.4095 | 1.4870 | 0.3289 | 1.9185 |
| Lak | Razorback_Lake | 42 | 179 | 0.1015 | 0.3945 | 1.6875 | 0.3233 | 1.9132 |
| Lak | Burntbridge_lake | 53 | 183 | 0.0651 | 0.3144 | 1.4920 | 0.4176 | 1.5863 |
| Lak | Gull_lake | 45 | 212 | 0.1047 | 0.3556 | 1.7135 | 0.2466 | 1.9865 |
| Lak | Alford_lake | 56 | 220 | 0.0702 | 0.3601 | 1.6862 | 0.2918 | 2.4132 |
| Lak | Safford_Lake | 44 | 225 | 0.1162 | 0.4022 | 1.7549 | 0.2759 | 2.0719 |
| Lak | Little_Rainbow_Lake | 52 | 247 | 0.0913 | 0.2850 | 1.6843 | 0.2011 | 1.8721 |
| Lak | Horseshoe_Lake | 49 | 255 | 0.1062 | 0.4321 | 1.8225 | 0.3715 | 2.4551 |
| Lak | Balsam_lake | 50 | 261 | 0.1044 | 0.4118 | 1.5782 | 0.3186 | 2.4367 |
| Lak | Beaver_lake | 56 | 267 | 0.0851 | 0.4260 | 1.6498 | 0.3591 | 2.4421 |
| Lak | Rat_Lake | 50 | 273 | 0.1092 | 0.4439 | 1.7768 | 0.2902 | 2.4133 |
| Lak | Stink_Lake | 53 | 281 | 0.1000 | 0.3520 | 1.7028 | 0.3394 | 2.5287 |
| Lak | Big_hope_lake | 61 | 328 | 0.0881 | 0.3900 | 1.7390 | 0.3733 | 2.1956 |
| Lak | Chub_pond | 54 | 416 | 0.1427 | 0.4217 | 1.9247 | 0.1665 | 3.4521 |
| Lak | Long_Lake | 65 | 417 | 0.0987 | 0.4685 | 1.7713 | 0.2763 | 3.5007 |
| Lak | Connera_lake | 65 | 418 | 0.0989 | 0.4005 | 1.6924 | 0.3871 | 2.9254 |
| Lak | Bridge_brook_lake | 75 | 553 | 0.0983 | 0.3748 | 1.6389 | 0.3702 | 3.6464 |
| Lak | Hoel_lake | 72 | 571 | 0.1101 | 0.4130 | 1.8399 | 0.3086 | 3.4722 |
| Lak | Meta | 211 | 8426 | 0.1893 | 3.3100 | 1.6677 | 0.1549 | 4.7178 |

Table S4: Empirical local food web values and 99% CIs for network properties: number of species $S$, connectance $C$, mean trophic level $TL$, where 99% CIs are based on 1000 networks generated by the assembly model.

| Metaweb | Network | $S$ | $S_{low}$ | $S_{high}$ | $C$ | $C_{low}$ | $C_{high}$ | $TL$ | $TL_{low}$ | $TL_{high}$ |
|---|---|---|---|---|---|---|---|---|---|---|
| Ant | Potter | 92 | 53.000 | 168.005 | 0.0383 | 0.0157 | 0.0714 | 2.1414 | 1.6182 | 2.4766 |
| Ant | Weddell | 435 | 339.975 | 511.005 | 0.0104 | 0.0099 | 0.0184 | 2.7908* | 1.8504 | 2.3804 |
| Isl | E1 | 25 | 10.995 | 44.005 | 0.1824 | 0.1790 | 0.5100 | 2.1108* | 2.2395 | 14.6920 |
| Isl | E2 | 31 | 18.000 | 60.005 | 0.2914 | 0.1712 | 0.4691 | 2.6022 | 2.2122 | 14.0229 |
| Isl | E3 | 24 | 13.000 | 46.010 | 0.2569 | 0.1662 | 0.4896 | 2.3143 | 1.8917 | 16.9056 |
| Isl | E7 | 24 | 13.000 | 46.000 | 0.2049 | 0.1575 | 0.4649 | 2.2584 | 1.9366 | 11.0201 |
| Isl | E9 | 33 | 28.000 | 73.000 | 0.2057 | 0.1696 | 0.4170 | 2.8061 | 2.3281 | 8.9635 |
| Isl | ST2 | 30 | 19.000 | 60.005 | 0.2311 | 0.1758 | 0.4500 | 2.3627 | 2.2822 | 10.5613 |
| Lak | Alford_lake | 56 | 26.990 | 81.005 | 0.0701 | 0.0354 | 0.1289 | 1.6861 | 1.3312 | 2.0473 |
| Lak | Balsam_lake | 50 | 25.985 | 78.000 | 0.1044 | 0.0337 | 0.1376 | 1.5782 | 1.2930 | 2.0869 |

| Metaweb | Network | $S$ | $S_{low}$ | $S_{high}$ | $C$ | $C_{low}$ | $C_{high}$ | $TL$ | $TL_{low}$ | $TL_{high}$ |
|---|---|---|---|---|---|---|---|---|---|---|
| Lak | Beaver_lake | 56 | 20.990 | 75.005 | 0.0851 | 0.0329 | 0.1355 | 1.6498 | 1.2753 | 2.0905 |
| Lak | Big_hope_lake | 61 | 16.980 | 74.005 | 0.0881 | 0.0349 | 0.1479 | 1.7390 | 1.2488 | 2.1694 |
| Lak | Brandy_lake | 30 | 7.995 | 53.015 | 0.1344 | 0.0377 | 0.2200 | 1.9132 | 1.1576 | 2.2326 |
| Lak | Bridge_brook_lake | 75 | 48.995 | 103.000 | 0.0983* | 0.0339 | 0.0825 | 1.6388 | 1.3173 | 1.9404 |
| Lak | Brook_trout_lake | 15 | 2.000 | 33.000 | 0.0844 | 0.0452 | 0.4286 | 1.5111 | 1.0908 | 2.2593 |
| Lak | Buck_pond | 41 | 13.995 | 63.000 | 0.0910 | 0.0372 | 0.1824 | 1.5300 | 1.1973 | 2.1673 |
| Lak | Burntbridge_lake | 53 | 19.000 | 71.000 | 0.0651 | 0.0346 | 0.1496 | 1.4920 | 1.2457 | 2.2135 |
| Lak | Cascade_lake | 35 | 12.995 | 63.000 | 0.0963 | 0.0366 | 0.1937 | 1.6399 | 1.2352 | 2.2246 |
| Lak | Chub_lake | 36 | 10.995 | 56.005 | 0.0640 | 0.0363 | 0.1625 | 1.4614 | 1.1763 | 2.1540 |
| Lak | Chub_pond | 54 | 25.000 | 78.005 | 0.1426* | 0.0330 | 0.1193 | 1.9247 | 1.2801 | 2.0316 |
| Lak | Connera_lake | 65 | 29.995 | 85.010 | 0.0989 | 0.0343 | 0.1095 | 1.6924 | 1.3284 | 2.0172 |
| Lak | Constable_lake | 32 | 9.000 | 55.000 | 0.0595 | 0.0382 | 0.2200 | 1.4107 | 1.1499 | 2.2153 |
| Lak | Deep_lake | 19 | 2.000 | 38.000 | 0.0775 | 0.0411 | 0.5000 | 1.3976 | 1.0908 | 2.4574 |
| Lak | Emerald_lake | 22 | 2.000 | 39.005 | 0.1198 | 0.0362 | 0.3601 | 1.6363 | 1.0587 | 2.4061 |
| Lak | Falls_lake | 39 | 13.995 | 62.005 | 0.0999 | 0.0350 | 0.1856 | 1.6161 | 1.2083 | 2.1627 |
| Lak | Fawn_lake | 32 | 7.995 | 56.000 | 0.1191 | 0.0356 | 0.2000 | 1.7077 | 1.1666 | 2.2092 |
| Lak | Federation_lake | 22 | 1.000 | 40.005 | 0.1177 | 0.0000 | 0.3333 | 1.7339 | 1.0000 | 2.4278 |
| Lak | Goose_lake | 40 | 15.000 | 61.000 | 0.0612 | 0.0340 | 0.1543 | 1.4657 | 1.1814 | 2.0956 |
| Lak | Grass_lake | 44 | 17.995 | 70.005 | 0.0852 | 0.0365 | 0.1412 | 1.6422 | 1.2996 | 2.1481 |
| Lak | Gull_lake | 45 | 23.000 | 71.005 | 0.1046 | 0.0353 | 0.1403 | 1.7135 | 1.2528 | 2.1865 |
| Lak | Gull_lake_north | 16 | 2.000 | 34.000 | 0.0976 | 0.0454 | 0.5000 | 1.4732 | 1.0869 | 2.3125 |
| Lak | Helldiver_pond | 41 | 12.990 | 63.000 | 0.1005 | 0.0353 | 0.1701 | 1.5881 | 1.2209 | 2.1819 |
| Lak | High_pond | 24 | 3.000 | 42.005 | 0.1510 | 0.0404 | 0.3333 | 1.8835 | 1.0999 | 2.5004 |
| Lak | Hoel_lake | 72 | 34.980 | 87.000 | 0.1101 | 0.0352 | 0.1158 | 1.8399 | 1.3094 | 2.0960 |
| Lak | Horseshoe_Lake | 49 | 23.000 | 78.010 | 0.1062 | 0.0315 | 0.1296 | 1.8224 | 1.2894 | 2.1229 |
| Lak | Indian_Lake | 35 | 8.995 | 51.000 | 0.0832 | 0.0366 | 0.2133 | 1.4590 | 1.1817 | 2.2074 |
| Lak | Little_Rainbow_Lake | 52 | 26.000 | 80.000 | 0.0913 | 0.0356 | 0.1332 | 1.6843 | 1.2939 | 2.0364 |
| Lak | Long_Lake | 65 | 31.995 | 86.005 | 0.0986 | 0.0345 | 0.1045 | 1.7713 | 1.3364 | 2.0960 |
| Lak | Loon_Lake | 35 | 11.995 | 62.000 | 0.0963 | 0.0347 | 0.1944 | 1.6612 | 1.1578 | 2.3640 |
| Lak | Lost_Lake | 31 | 7.990 | 54.005 | 0.1540 | 0.0386 | 0.2400 | 1.8109 | 1.1630 | 2.3057 |
| Lak | Lost_Lake_East | 41 | 10.990 | 57.025 | 0.0814 | 0.0366 | 0.1769 | 1.5691 | 1.1904 | 2.2217 |
| Lak | Lower_Sister_Lake | 37 | 13.995 | 63.005 | 0.1176 | 0.0334 | 0.1901 | 1.7332 | 1.1773 | 2.2070 |
| Lak | Oswego_Lake | 33 | 8.995 | 54.005 | 0.1267 | 0.0381 | 0.2100 | 1.6789 | 1.1175 | 2.2125 |
| Lak | Owl_Lake | 30 | 6.000 | 48.000 | 0.0788 | 0.0394 | 0.2500 | 1.4212 | 1.1666 | 2.3150 |
| Lak | Rat_Lake | 50 | 26.985 | 80.000 | 0.1092 | 0.0335 | 0.1234 | 1.7767 | 1.3029 | 2.0791 |
| Lak | Razorback_Lake | 42 | 11.995 | 61.005 | 0.1014 | 0.0335 | 0.1611 | 1.6875 | 1.1874 | 2.1844 |
| Lak | Rock_Lake | 22 | 3.995 | 43.005 | 0.0888 | 0.0425 | 0.3125 | 1.6201 | 1.0833 | 2.4286 |
| Lak | Russian_Lake | 24 | 6.990 | 48.000 | 0.1059 | 0.0400 | 0.2231 | 1.6815 | 1.0909 | 2.2801 |
| Lak | Safford_Lake | 44 | 20.000 | 73.005 | 0.1162 | 0.0344 | 0.1384 | 1.7549 | 1.2662 | 2.1500 |
| Lak | Sand_Lake | 29 | 6.000 | 46.015 | 0.0975 | 0.0384 | 0.2422 | 1.6248 | 1.1110 | 2.2941 |
| Lak | South_Lake | 22 | 2.000 | 37.000 | 0.0743 | 0.0399 | 0.4075 | 1.4090 | 1.0833 | 2.2795 |
| Lak | Squaw_Lake | 41 | 12.000 | 63.005 | 0.1005 | 0.0367 | 0.1737 | 1.4869 | 1.1814 | 2.2544 |
| Lak | Stink_Lake | 53 | 26.000 | 79.000 | 0.1000 | 0.0365 | 0.1291 | 1.7028 | 1.2808 | 2.1233 |
| Lak | Twelfth_Tee_Lake | 31 | 6.000 | 50.005 | 0.0863 | 0.0375 | 0.2500 | 1.5079 | 1.1325 | 2.2212 |
| Lak | Twin_Lake_East | 13 | 1.000 | 25.005 | 0.1005 | 0.0000 | 0.5628 | 1.4871 | 1.0000 | 2.3750 |
| Lak | Twin_Lake_West | 26 | 7.995 | 55.005 | 0.0887 | 0.0367 | 0.2231 | 1.4846 | 1.1537 | 2.2325 |
| Lak | Whipple_Lake | 32 | 8.000 | 55.000 | 0.1328 | 0.0378 | 0.2187 | 1.9117 | 1.1332 | 2.2327 |
| Lak | Wolf_Lake | 27 | 4.990 | 45.000 | 0.0576 | 0.0399 | 0.3087 | 1.2798 | 1.1248 | 2.3540 |

Table S5: Empirical local food web values and 99% CIs for network properties: trophic coherence $Q$, modularity $MO$, and mean maximal eingenvalue $MEing$, where 99% CIs are based on 1000 networks generated by the assembly model.

| Metaweb | Network | $Q$ | $Q_{low}$ | $Q_{high}$ | $MO$ | $MO_{low}$ | $MO_{high}$ | $MEing$ | $MEing_{low}$ | $MEing_{high}$ |
|---|---|---|---|---|---|---|---|---|---|---|
| Ant | Potter | 0.5230 | 0.3019 | 0.8918 | 0.3640 | 0.3118 | 0.6339 | 2.1620 | 0.4407 | 3.8803 |
| Ant | Weddell | 0.5829 | 0.5670 | 0.8380 | 0.4585 | 0.4204 | 0.5507 | 3.6264* | 3.6833 | 5.9177 |
| Isl | E1 | 0.5501* | 0.5965 | 2.6662 | 0.1721 | 0.1129 | 0.2620 | 2.0248 | 1.3927 | 8.2489 |
| Isl | E2 | 0.7507 | 0.6343 | 2.5077 | 0.1601 | 0.1144 | 0.2408 | 4.0469 | 2.8248 | 9.4923 |
| Isl | E3 | 0.6157 | 0.5963 | 3.2707 | 0.1334 | 0.1147 | 0.2616 | 2.9558 | 1.8152 | 8.3739 |
| Isl | E7 | 0.7047 | 0.5599 | 2.4501 | 0.1765 | 0.1100 | 0.2762 | 2.8429 | 1.6886 | 8.6337 |
| Isl | E9 | 0.7660 | 0.6674 | 2.3052 | 0.2437* | 0.1153 | 0.2232 | 5.3367 | 4.7997 | 10.4595 |
| Isl | ST2 | 0.7006 | 0.6484 | 2.5031 | 0.1561 | 0.1147 | 0.2385 | 3.6717 | 3.0549 | 9.2790 |
| Lak | Alford_lake | 0.3601 | 0.2169 | 0.6585 | 0.2909 | 0.1708 | 0.4529 | 2.4132 | 0.0830 | 3.0660 |
| Lak | Balsam_lake | 0.4118 | 0.1981 | 0.6184 | 0.3174 | 0.1897 | 0.4592 | 2.4367 | 0.2034 | 2.8791 |
| Lak | Beaver_lake | 0.4260 | 0.2306 | 0.6756 | 0.3578 | 0.1859 | 0.4806 | 2.4421 | -0.0085 | 3.0042 |
| Lak | Big_hope_lake | 0.3900 | 0.1709 | 0.6692 | 0.3718 | 0.1671 | 0.4846 | 2.1956 | -0.0462 | 2.7280 |
| Lak | Brandy_lake | 0.4179 | 0.0000 | 0.7280 | 0.1198 | 0.0740 | 0.4849 | 1.7252 | -0.1697 | 2.3065 |
| Lak | Bridge_brook_lake | 0.3748 | 0.2518 | 0.5836 | 0.3693 | 0.2257 | 0.4395 | 3.6464* | 0.6357 | 3.4227 |
| Lak | Brook_trout_lake | 0.3244 | 0.0000 | 0.8165 | 0.2906 | -0.0050 | 0.4765 | -0.1133 | -0.4792 | 1.7764 |
| Lak | Buck_pond | 0.1974 | 0.0000 | 0.6726 | 0.3836 | 0.1583 | 0.4802 | 0.8899 | -0.0782 | 2.6142 |
| Lak | Burntbridge_lake | 0.3144 | 0.1834 | 0.6802 | 0.4174 | 0.1730 | 0.4844 | 1.5863 | -0.0255 | 2.8088 |
| Lak | Cascade_lake | 0.3983 | 0.1549 | 0.7180 | 0.3775 | 0.1419 | 0.4956 | 1.3142 | -0.0666 | 2.5310 |
| Lak | Chub_lake | 0.4514 | 0.0000 | 0.6884 | 0.3047 | 0.1222 | 0.4884 | 1.1629 | -0.1297 | 2.4811 |
| Lak | Chub_pond | 0.4217 | 0.2057 | 0.6419 | 0.1660* | 0.1770 | 0.4777 | 3.4521* | 0.2089 | 2.8156 |
| Lak | Connera_lake | 0.4005 | 0.2380 | 0.6080 | 0.3857 | 0.1974 | 0.4590 | 2.9254 | 0.2675 | 3.2941 |
| Lak | Constable_lake | 0.4090 | 0.0000 | 0.7213 | 0.3668 | 0.1232 | 0.4813 | 0.9281 | -0.1881 | 2.1424 |
| Lak | Deep_lake | 0.3164 | 0.0000 | 0.8930 | 0.3670 | 0.0000 | 0.4752 | -0.0682 | -0.3889 | 1.8846 |
| Lak | Emerald_lake | 0.4828 | 0.0000 | 0.8734 | 0.2512 | -0.0041 | 0.4731 | 0.9974 | -0.4673 | 1.9050 |
| Lak | Falls_lake | 0.2954 | 0.0981 | 0.7092 | 0.3499 | 0.1632 | 0.4890 | 1.2022 | -0.1054 | 2.7316 |
| Lak | Fawn_lake | 0.3640 | 0.0000 | 0.7416 | 0.2738 | 0.1482 | 0.4759 | 1.4517 | -0.1457 | 2.2932 |
| Lak | Federation_lake | 0.3508 | 0.0000 | 0.8242 | 0.3192 | -0.0035 | 0.4810 | 0.5067 | -0.4801 | 1.8439 |
| Lak | Goose_lake | 0.4375 | 0.0000 | 0.6619 | 0.3346 | 0.1449 | 0.4964 | 1.5594 | -0.0882 | 2.6237 |
| Lak | Grass_lake | 0.3979 | 0.2392 | 0.6756 | 0.2949 | 0.1769 | 0.4764 | 1.6909 | -0.0387 | 2.8513 |
| Lak | Gull_lake | 0.3556 | 0.1808 | 0.6638 | 0.2486 | 0.1611 | 0.4669 | 1.9865 | -0.0269 | 2.8248 |
| Lak | Gull_lake_north | 0.4121 | 0.0000 | 0.8650 | 0.3010 | -0.0035 | 0.4663 | -0.0735 | -0.4803 | 1.5415 |
| Lak | Helldiver_pond | 0.3787 | 0.1706 | 0.6709 | 0.2445 | 0.1612 | 0.4786 | 1.3853 | -0.0715 | 2.6919 |
| Lak | High_pond | 0.4843 | 0.0000 | 0.8759 | 0.1893 | -0.0038 | 0.4694 | 1.3283 | -0.4614 | 1.9935 |
| Lak | Hoel_lake | 0.4130 | 0.2068 | 0.6052 | 0.3081 | 0.1961 | 0.4592 | 3.4722* | 0.1834 | 3.2154 |
| Lak | Horseshoe_Lake | 0.4321 | 0.1598 | 0.6179 | 0.3711 | 0.1811 | 0.4731 | 2.4551 | 0.0926 | 2.9783 |
| Lak | Indian_Lake | 0.2173 | 0.0000 | 0.7252 | 0.3913 | 0.1352 | 0.4834 | 0.2044 | -0.2328 | 2.2488 |
| Lak | Little_Rainbow_Lake | 0.2850 | 0.2234 | 0.6190 | 0.2010 | 0.1713 | 0.4671 | 1.8721 | 0.0201 | 3.1177 |
| Lak | Long_Lake | 0.4685 | 0.2331 | 0.6079 | 0.2756 | 0.2127 | 0.4581 | 3.5007* | 0.1407 | 3.0996 |
| Lak | Loon_Lake | 0.4436 | 0.0951 | 0.7511 | 0.1781 | 0.1423 | 0.4850 | 1.4555 | -0.0803 | 2.7344 |
| Lak | Lost_Lake | 0.3244 | 0.0000 | 0.7396 | 0.2505 | 0.0800 | 0.4723 | 1.1592 | -0.1395 | 2.3895 |
| Lak | Lost_Lake_East | 0.2734 | 0.0000 | 0.7111 | 0.4119 | 0.1414 | 0.4805 | 1.2536 | -0.1084 | 2.4022 |
| Lak | Lower_Sister_Lake | 0.4090 | 0.0000 | 0.7106 | 0.2438 | 0.1270 | 0.4834 | 1.9143 | -0.1059 | 2.5285 |
| Lak | Oswego_Lake | 0.3659 | 0.0000 | 0.7158 | 0.3229 | 0.0000 | 0.4680 | 1.2401 | -0.1959 | 2.2655 |
| Lak | Owl_Lake | 0.2010 | 0.0000 | 0.7701 | 0.3914 | 0.0559 | 0.4905 | 0.1351 | -0.1884 | 2.1956 |
| Lak | Rat_Lake | 0.4439 | 0.2235 | 0.6210 | 0.2911 | 0.1901 | 0.4766 | 2.4133 | 0.1787 | 3.0122 |
| Lak | Razorback_Lake | 0.3945 | 0.0000 | 0.6731 | 0.3222 | 0.1672 | 0.4751 | 1.9132 | -0.0774 | 2.6185 |

| Metaweb | Network | $Q$ | $Q_{low}$ | $Q_{high}$ | $MO$ | $MO_{low}$ | $MO_{high}$ | $MEing$ | $MEing_{low}$ | $MEing_{high}$ |
|---|---|---|---|---|---|---|---|---|---|---|
| Lak | Rock_Lake | 0.2859 | 0.0000 | 0.8102 | 0.2757 | -0.0026 | 0.4872 | 0.0069 | -0.4660 | 1.8356 |
| Lak | Russian_Lake | 0.3269 | 0.0000 | 0.7800 | 0.1993 | 0.0000 | 0.4887 | 0.8123 | -0.1916 | 2.0958 |
| Lak | Safford_Lake | 0.4022 | 0.2154 | 0.6741 | 0.2757 | 0.1943 | 0.4788 | 2.0719 | -0.0208 | 2.7132 |
| Lak | Sand_Lake | 0.4567 | 0.0000 | 0.8001 | 0.3382 | 0.0000 | 0.4892 | 1.0770 | -0.2446 | 2.0258 |
| Lak | South_Lake | 0.0000 | 0.0000 | 0.8465 | 0.3492 | -0.0038 | 0.4764 | 0.0417 | -0.4737 | 1.6940 |
| Lak | Squaw_Lake | 0.4095 | 0.0000 | 0.7299 | 0.3275 | 0.1156 | 0.4767 | 1.9185 | -0.0731 | 2.5158 |
| Lak | Stink_Lake | 0.3520 | 0.2177 | 0.6415 | 0.3313 | 0.1763 | 0.4729 | 2.5287 | 0.0424 | 2.9646 |
| Lak | Twelfth_Tee_Lake | 0.4122 | 0.0000 | 0.7861 | 0.3744 | 0.1088 | 0.4893 | 0.8816 | -0.2537 | 2.1222 |
| Lak | Twin_Lake_East | 0.3333 | 0.0000 | 0.9099 | 0.3937 | -0.0062 | 0.4505 | -0.1111 | -0.5097 | 1.3361 |
| Lak | Twin_Lake_West | 0.1653 | 0.0000 | 0.7467 | 0.3672 | 0.0948 | 0.4761 | 0.0968 | -0.1750 | 2.2301 |
| Lak | Whipple_Lake | 0.4370 | 0.0000 | 0.7145 | 0.1996 | 0.1272 | 0.4814 | 1.9576 | -0.1787 | 2.3403 |
| Lak | Wolf_Lake | 0.2583 | 0.0000 | 0.8266 | 0.3748 | 0.0000 | 0.4641 | 0.0100 | -0.2819 | 1.9356 |

Table S6: Motif counts for the empirical food webs and 99% CIs based on 1000 assembly model networks, where EC is exploitative competition, AC apparent competition, TT tri-trophic chain and OM is omnivory. Quantities marked with ’*’ are significant at 1% level. Only 1 network showed significant under-representation and 9 networks over-representation for at least one motif.

| Metaweb | Network | EC | $EC_{low}$ | $EC_{high}$ | AC | $AC_{low}$ | $AC_{high}$ | TT | $TT_{low}$ | $TT_{high}$ | OM | $OM_{low}$ | $OM_{high}$ |
|---|---|---|---|---|---|---|---|---|---|---|---|---|---|
| Ant | Potter | 973 | 297 | 7481 | 2123 | 191 | 2129 | 620 | 53 | 1153 | 139 | 3 | 750 |
| Ant | Weddell | 47573 | 21422 | 80409 | 7575* | 8577 | 24632 | 3846 | 3810 | 11999 | 1156* | 1509 | 7476 |
| Isl | E1 | 140 | 319 | 53 | 69 | 5 | 804 | 3 | 1528 | 2 | 555 | 2 | 805 |
| Isl | E2 | 372 | 783 | 163 | 300 | 33 | 1982 | 51 | 4173 | 19 | 1222 | 33 | 1708 |
| Isl | E3 | 184 | 418 | 96 | 150 | 13 | 1135 | 18 | 2550 | 6 | 733 | 9 | 975 |
| Isl | E7 | 100 | 199 | 58 | 54 | 10 | 1083 | 16 | 1942 | 7 | 676 | 9 | 907 |
| Isl | E9 | 377 | 506 | 293 | 180 | 207 | 3970 | 287 | 9734 | 143 | 2604 | 168 | 3865 |
| Isl | ST2 | 183 | 592 | 110 | 195 | 34 | 2116 | 65 | 4857 | 33 | 1277 | 44 | 1794 |
| Lak | Alford_lake | 360 | 19 | 1037 | 2059 | 176 | 3600 | 306 | 55 | 1525 | 170 | 0 | 433 |
| Lak | Balsam_lake | 445 | 26 | 855 | 2232 | 152 | 3112 | 403 | 39 | 1336 | 233 | 0 | 363 |
| Lak | Beaver_lake | 391 | 10 | 857 | 2011 | 66 | 2788 | 765 | 24 | 1229 | 235 | 0 | 329 |
| Lak | Big_hope_lake | 693* | 7 | 678 | 2747* | 45 | 2491 | 858 | 13 | 1149 | 258 | 0 | 273 |
| Lak | Brandy_lake | 174 | 0 | 324 | 813 | 6 | 1192 | 85 | 0 | 491 | 97 | 0 | 129 |
| Lak | Bridge_brook_lake | 1539 | 119 | 1663 | 6338* | 502 | 6036 | 1931 | 195 | 2628 | 629 | 15 | 762 |
| Lak | Brook_trout_lake | 15 | 0 | 96 | 46 | 0 | 302 | 10 | 0 | 152 | 0 | 0 | 40 |
| Lak | Buck_pond | 443 | 7 | 512 | 1051 | 27 | 1802 | 122 | 6 | 751 | 0 | 0 | 203 |
| Lak | Burntbridge_lake | 295 | 8 | 658 | 1599 | 58 | 2457 | 247 | 17 | 1049 | 55 | 0 | 279 |
| Lak | Cascade_lake | 126 | 3 | 554 | 539 | 15 | 1895 | 208 | 5 | 798 | 54 | 0 | 240 |
| Lak | Chub_lake | 72 | 2 | 360 | 408 | 18 | 1375 | 118 | 2 | 591 | 27 | 0 | 159 |
| Lak | Chub_pond | 1180* | 23 | 904 | 3100 | 144 | 3341 | 1048 | 48 | 1407 | 802* | 0 | 422 |
| Lak | Connera_lake | 787 | 23 | 1122 | 4035* | 206 | 3820 | 1455 | 82 | 1753 | 384 | 0 | 408 |
| Lak | Constable_lake | 72 | 0 | 323 | 255 | 5 | 1017 | 56 | 0 | 562 | 0 | 0 | 159 |
| Lak | Deep_lake | 21 | 0 | 168 | 70 | 0 | 550 | 19 | 0 | 258 | 0 | 0 | 67 |
| Lak | Emerald_lake | 48 | 0 | 172 | 136 | 0 | 650 | 70 | 0 | 247 | 0 | 0 | 75 |
| Lak | Falls_lake | 327 | 5 | 463 | 876 | 27 | 1635 | 268 | 5 | 788 | 40 | 0 | 214 |
| Lak | Fawn_lake | 165 | 0 | 351 | 512 | 6 | 1262 | 233 | 0 | 555 | 64 | 0 | 153 |
| Lak | Federation_lake | 85 | 0 | 156 | 171 | 0 | 488 | 64 | 0 | 263 | 9 | 0 | 60 |
| Lak | Goose_lake | 61 | 3 | 450 | 522 | 43 | 1568 | 151 | 2 | 667 | 57 | 0 | 210 |
| Lak | Grass_lake | 265 | 9 | 779 | 1027 | 44 | 2505 | 292 | 29 | 1181 | 119 | 0 | 299 |

| Metaweb | Network | EC | $EC_{low}$ | $EC_{high}$ | AC | $AC_{low}$ | $AC_{high}$ | TT | $TT_{low}$ | $TT_{high}$ | OM | $OM_{low}$ | $OM_{high}$ |
|---|---|---|---|---|---|---|---|---|---|---|---|---|---|
| Lak | Gull_lake | 514 | 8 | 751 | 1172 | 57 | 2678 | 434 | 17 | 1298 | 183 | 0 | 329 |
| Lak | Gull_lake_north | 21 | 0 | 91 | 47 | 0 | 313 | 18 | 0 | 143 | 0 | 0 | 40 |
| Lak | Helldiver_pond | 360 | 4 | 535 | 967 | 11 | 1758 | 354 | 0 | 733 | 92 | 0 | 205 |
| Lak | High_pond | 117 | 0 | 214 | 276 | 0 | 643 | 103 | 0 | 333 | 30 | 0 | 86 |
| Lak | Hoel_lake | 1393* | 47 | 1204 | 5109* | 211 | 4459 | 2488* | 70 | 1938 | 991* | 3 | 496 |
| Lak | Horseshoe_Lake | 352 | 26 | 1010 | 1595 | 126 | 3285 | 699 | 49 | 1373 | 213 | 0 | 388 |
| Lak | Indian_Lake | 281* | 0 | 276 | 581 | 3 | 937 | 72 | 0 | 483 | 0 | 0 | 134 |
| Lak | Little_Rainbow_Lake | 915* | 22 | 873 | 2094 | 89 | 3280 | 467 | 38 | 1318 | 175 | 0 | 351 |
| Lak | Long_Lake | 827 | 34 | 1011 | 3756 | 181 | 3881 | 1237 | 59 | 1667 | 656* | 0 | 430 |
| Lak | Loon_Lake | 159 | 2 | 471 | 635 | 20 | 1800 | 190 | 0 | 700 | 81 | 0 | 189 |
| Lak | Lost_Lake | 327 | 0 | 402 | 623 | 5 | 1197 | 348 | 0 | 548 | 48 | 0 | 141 |
| Lak | Lost_Lake_East | 265 | 1 | 404 | 893 | 8 | 1408 | 181 | 0 | 610 | 35 | 0 | 158 |
| Lak | Lower_Sister_Lake | 249 | 2 | 480 | 880 | 21 | 1611 | 244 | 4 | 763 | 149 | 0 | 216 |
| Lak | Oswego_Lake | 256 | 0 | 341 | 608 | 5 | 1232 | 250 | 0 | 548 | 30 | 0 | 144 |
| Lak | Owl_Lake | 129 | 0 | 286 | 367 | 2 | 910 | 29 | 0 | 450 | 0 | 0 | 104 |
| Lak | Rat_Lake | 398 | 20 | 943 | 1675 | 138 | 3175 | 721 | 34 | 1361 | 259 | 0 | 363 |
| Lak | Razorback_Lake | 291 | 4 | 500 | 1048 | 15 | 1750 | 298 | 6 | 804 | 96 | 0 | 210 |
| Lak | Rock_Lake | 113 | 0 | 155 | 107 | 1 | 593 | 22 | 0 | 276 | 0 | 0 | 68 |
| Lak | Russian_Lake | 110 | 0 | 263 | 212 | 2 | 979 | 64 | 0 | 474 | 20 | 0 | 111 |
| Lak | Safford_Lake | 406 | 9 | 694 | 1301 | 52 | 2521 | 564 | 2 | 1076 | 206 | 0 | 244 |
| Lak | Sand_Lake | 68 | 0 | 276 | 277 | 0 | 982 | 133 | 0 | 402 | 28 | 0 | 124 |
| Lak | South_Lake | 100 | 0 | 150 | 90 | 0 | 482 | 0 | 0 | 214 | 0 | 0 | 60 |
| Lak | Squaw_Lake | 234 | 4 | 526 | 1197 | 23 | 1783 | 303 | 10 | 754 | 88 | 0 | 202 |
| Lak | Stink_Lake | 596 | 22 | 935 | 2278 | 114 | 3246 | 656 | 47 | 1474 | 221 | 0 | 388 |
| Lak | Twelfth_Tee_Lake | 89 | 0 | 265 | 318 | 6 | 822 | 96 | 0 | 353 | 16 | 0 | 107 |
| Lak | Twin_Lake_East | 16 | 0 | 69 | 20 | 0 | 193 | 4 | 0 | 91 | 0 | 0 | 20 |
| Lak | Twin_Lake_West | 127 | 1 | 394 | 219 | 4 | 1364 | 11 | 0 | 553 | 0 | 0 | 178 |
| Lak | Whipple_Lake | 199 | 0 | 380 | 582 | 8 | 1262 | 200 | 0 | 579 | 154 | 0 | 167 |
| Lak | Wolf_Lake | 45 | 0 | 241 | 182 | 0 | 806 | 33 | 0 | 368 | 0 | 0 | 76 |

Table S7: Comparison of empirical proportions of topological roles with one assembly model realization corresponding to Figure 6, run 19 of table S8. In this case only 10% of the total (6/58), are significantly different from the metaweb assembly model at p=0.01 level.

| Metaweb | Network | Size | chi2 | pvalue |
|---|---|---|---|---|
| Ant | Weddell | 435 | 10.3035 | 0.0154 |
| Ant | Potter * | 92 | 31.6872 | 0.0001 |
| Lak | Bridge brook lake | 75 | 3.4161 | 0.2468 |
| Lak | Hoel lake | 72 | 5.3088 | 0.0336 |
| Lak | Connera lake | 65 | 1.8070 | 0.4988 |
| Lak | Long Lake | 65 | 2.6933 | 0.2448 |
| Lak | Big hope lake * | 61 | 8.2453 | 0.0066 |
| Lak | Alford lake | 56 | 5.8061 | 0.0274 |
| Lak | Beaver lake | 56 | 0.0043 | 1.0000 |
| Lak | Chub pond | 54 | 2.5568 | 0.2533 |
| Lak | Burntbridge lake | 53 | 1.0865 | 1.0000 |
| Lak | Stink Lake | 53 | 0.0050 | 1.0000 |
| Lak | Little Rainbow Lake | 52 | 3.6207 | 0.4073 |

| Metaweb | Network | Size | chi2 | pvalue |
|---|---|---|---|---|
| Lak | Balsam lake | 50 | 2.3808 | 0.3181 |
| Lak | Rat Lake | 50 | 1.6422 | 1.0000 |
| Lak | Horseshoe Lake | 49 | 2.0846 | 0.2474 |
| Lak | Gull lake | 45 | 0.8418 | 1.0000 |
| Lak | Grass lake | 44 | 0.6712 | 0.5803 |
| Lak | Safford Lake | 44 | 1.4234 | 1.0000 |
| Lak | Razorback Lake | 42 | 1.0630 | 0.4915 |
| Lak | Buck pond | 41 | 2.2905 | 0.2155 |
| Lak | Helldiver pond | 41 | 2.2745 | 0.2324 |
| Lak | Lost Lake East * | 41 | 7.4108 | 0.0091 |
| Lak | Squaw Lake | 41 | 1.0646 | 0.4853 |
| Lak | Goose lake | 40 | 0.3210 | 1.0000 |
| Lak | Falls lake | 39 | 1.6289 | 1.0000 |
| Lak | Lower Sister Lake | 37 | 1.9420 | 0.5159 |
| Lak | Chub lake | 36 | 7.7269 | 0.0103 |
| Lak | Cascade lake | 35 | 3.0000 | 0.2517 |
| Lak | Indian Lake | 35 | 3.1660 | 0.4562 |
| Lak | Loon Lake | 35 | 4.5957 | 0.0901 |
| Lak | Oswego Lake | 33 | 4.9993 | 0.1499 |
| Lak | Constable lake | 32 | 0.9161 | 1.0000 |
| Lak | Fawn lake | 32 | 4.0809 | 0.1646 |
| Lak | Whipple Lake | 32 | 0.9860 | 1.0000 |
| Lak | Lost Lake | 31 | 0.5501 | 1.0000 |
| Lak | Twelfth Tee Lake | 31 | 3.6631 | 0.2399 |
| Lak | Brandy lake | 30 | 1.0548 | 0.7450 |
| Lak | Owl Lake | 30 | 4.0127 | 0.1167 |
| Lak | Sand Lake | 29 | 3.2464 | 0.5009 |
| Lak | Wolf Lake | 27 | 0.8318 | 1.0000 |
| Lak | Twin Lake West | 26 | 1.1540 | 0.4867 |
| Lak | High pond | 24 | 2.0017 | 1.0000 |
| Lak | Russian Lake | 24 | 1.5074 | 1.0000 |
| Lak | Emerald lake | 22 | 3.6808 | 0.4125 |
| Lak | Federation lake | 22 | 0.9370 | 1.0000 |
| Lak | Rock Lake | 22 | 4.1000 | 0.2333 |
| Lak | South Lake | 22 | 0.0219 | 1.0000 |
| Lak | Deep lake | 19 | 1.4714 | 0.4967 |
| Lak | Gull lake north | 16 | 0.6748 | 1.0000 |
| Lak | Brook trout lake | 15 | 2.9156 | 0.1389 |
| Lak | Twin Lake East | 13 | 1.1285 | 0.4802 |
| Isl | E9 | 33 | 3.8293 | 0.0693 |
| Isl | E2 * | 31 | 13.3309 | 0.0002 |
| Isl | ST2 * | 30 | 7.8048 | 0.0075 |
| Isl | E1 | 25 | 0.1963 | 1.0000 |
| Isl | E7 * | 24 | 9.7222 | 0.0041 |
| Isl | E3 | 24 | 2.1797 | 0.2304 |

Table S8: Comparison of empirical proportions of topological roles with 20 different assembly model realizations at 0.01 significance level. The number of significant results varies between 3 and 10, which represents betwenn 5% and 17% significant results.

| run | Number of significant | total | Freq |
|---:|---:|---:|---:|
| 0 | 6 | 58 | 0.10 |
| 1 | 5 | 58 | 0.09 |
| 2 | 8 | 58 | 0.14 |
| 3 | 7 | 58 | 0.12 |
| 4 | 6 | 58 | 0.10 |
| 5 | 3 | 58 | 0.05 |
| 6 | 3 | 58 | 0.05 |
| 7 | 6 | 58 | 0.10 |
| 8 | 9 | 58 | 0.16 |
| 9 | 5 | 58 | 0.09 |
| 10 | 8 | 58 | 0.14 |
| 11 | 6 | 58 | 0.10 |
| 12 | 4 | 58 | 0.07 |
| 13 | 5 | 58 | 0.09 |
| 14 | 7 | 58 | 0.12 |
| 15 | 7 | 58 | 0.12 |
| 16 | 8 | 58 | 0.14 |
| 17 | 10 | 58 | 0.17 |
| 18 | 4 | 58 | 0.07 |
| 19 | 6 | 58 | 0.10 |

Table S9: Comparison of empirical proportions of topological roles with 20 different assembly model realizations at 0.01 significance level. Only Potter Cove is different in 100% of the comparisons, Island E9 is different 60% of the times, and the other food webs showed differences less often.

| meta | Network | significant | total | Freq |
|---|---|---:|---:|---:|
| Ant | Potter | 20 | 20 | 1.00 |
| Ant | Weddell | 10 | 20 | 0.50 |
| Isl | E1 | 3 | 20 | 0.15 |
| Isl | E2 | 11 | 20 | 0.55 |
| Isl | E3 | 10 | 20 | 0.50 |
| Isl | E7 | 8 | 20 | 0.40 |
| Isl | E9 | 12 | 20 | 0.60 |
| Isl | ST2 | 5 | 20 | 0.25 |
| Lak | Alford lake | 1 | 20 | 0.05 |
| Lak | Beaver lake | 1 | 20 | 0.05 |
| Lak | Big hope lake | 4 | 20 | 0.20 |
| Lak | Brandy lake | 1 | 20 | 0.05 |
| Lak | Bridge brook lake | 3 | 20 | 0.15 |
| Lak | Buck pond | 1 | 20 | 0.05 |
| Lak | Burntbridge lake | 2 | 20 | 0.10 |
| Lak | Cascade lake | 1 | 20 | 0.05 |
| Lak | Chub lake | 2 | 20 | 0.10 |
| Lak | Chub pond | 1 | 20 | 0.05 |
| Lak | Connera lake | 1 | 20 | 0.05 |

| meta | Network | significant | total | Freq |
|---|---|---|---|---|
| Lak | Federation lake | 1 | 20 | 0.05 |
| Lak | Goose lake | 1 | 20 | 0.05 |
| Lak | Gull lake | 3 | 20 | 0.15 |
| Lak | Helldiver pond | 1 | 20 | 0.05 |
| Lak | Hoel lake | 1 | 20 | 0.05 |
| Lak | Horseshoe Lake | 1 | 20 | 0.05 |
| Lak | Little Rainbow Lake | 1 | 20 | 0.05 |
| Lak | Long Lake | 2 | 20 | 0.10 |
| Lak | Loon Lake | 1 | 20 | 0.05 |
| Lak | Lost Lake | 1 | 20 | 0.05 |
| Lak | Lost Lake East | 3 | 20 | 0.15 |
| Lak | Oswego Lake | 2 | 20 | 0.10 |
| Lak | Razorback Lake | 2 | 20 | 0.10 |
| Lak | Sand Lake | 1 | 20 | 0.05 |
| Lak | Stink Lake | 1 | 20 | 0.05 |
| Lak | Twelfth Tee Lake | 1 | 20 | 0.05 |
| Lak | Twin Lake West | 2 | 20 | 0.10 |
| Lak | Wolf Lake | 1 | 20 | 0.05 |

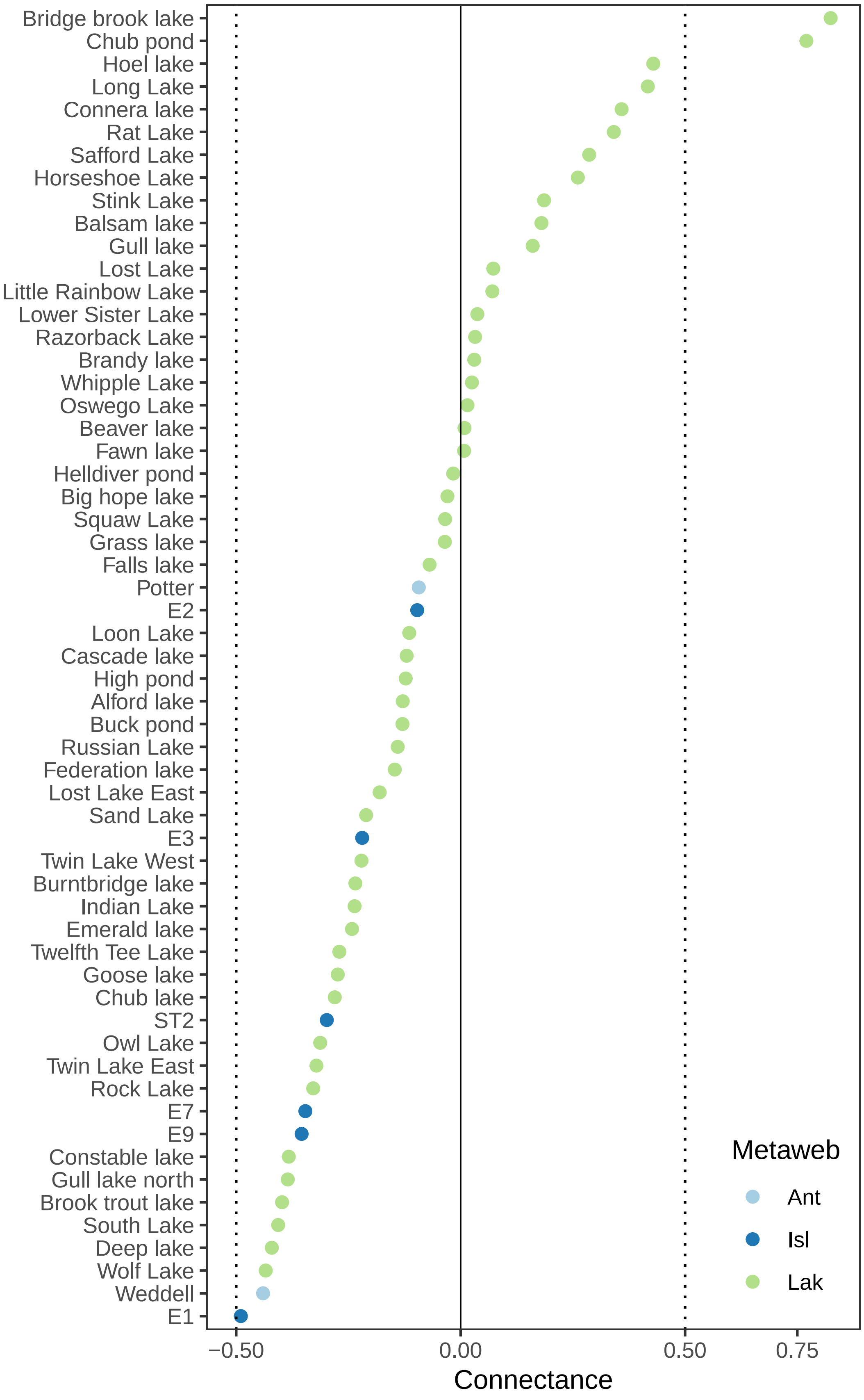


Figure S3: Connectance comparison for local empirical networks (dots) and assembly null model networks. We ran 1000 simulations of the model fitted to local networks to build the 99% CIs of the network metric (vertical dotted lines). Colors represent metawebs to which local food webs belong, where *Ant* is the Antarctic, *Isl* the Islands, and *Lak* the lakes metawebs.

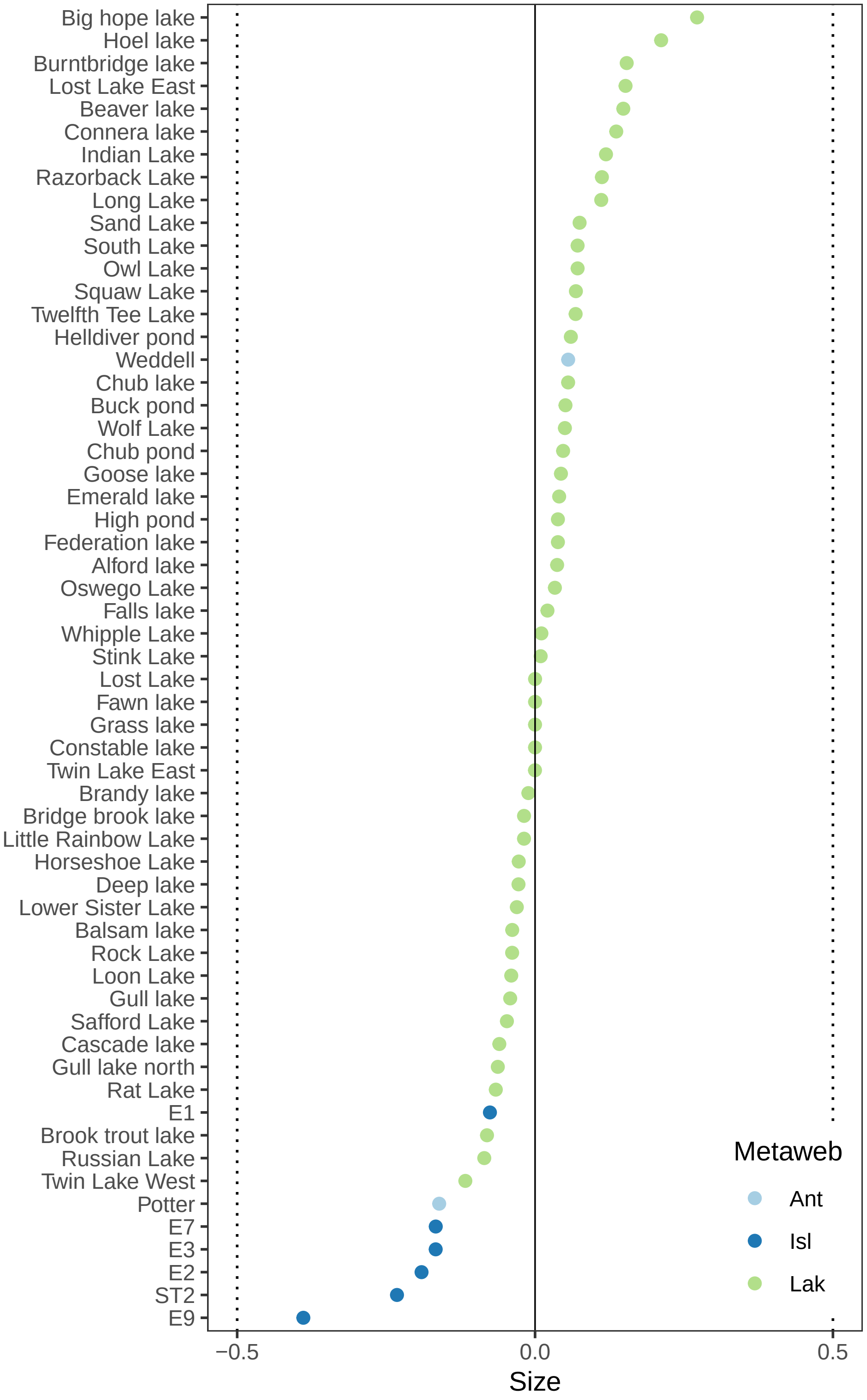


Figure S4: Number of species (Size) comparison for local empirical networks (dots) and assembly null model networks. We ran 1000 simulations of the model fitted to local networks to build the 99% CIs of the network metric (vertical dotted lines). Colors represent metawebs to which local food webs belong, where *Ant* is the Antarctic, *Isl* the Islands, and *Lak* the lakes metawebs.

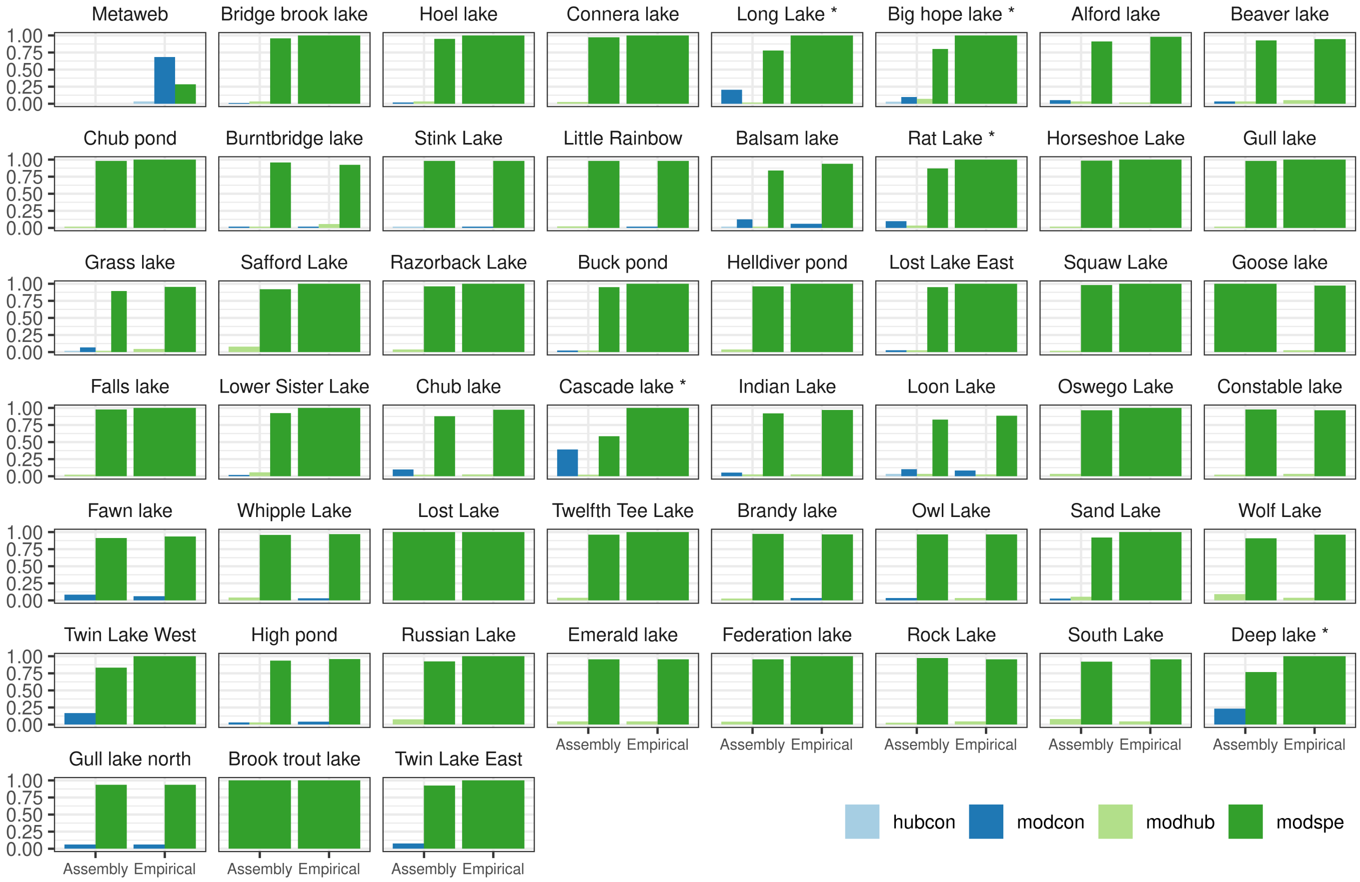


Figure S5: Topological roles comparison for local empirical networks (dots) and assembly model networks for the Lakes metaweb. Only 3 local food webs have proportions different from the metaweb assembly model. The topological roles are: *Hub connectors* (hubcon) have a high number of between module links, *Module connectors* (modcon) have a low number of links mostly between modules, *Module hubs* (modhub) have a high number of links inside its module, *Module specialists* (modspe) have a low number of links inside its module. Lakes marked with '*' are significant at 1% level.

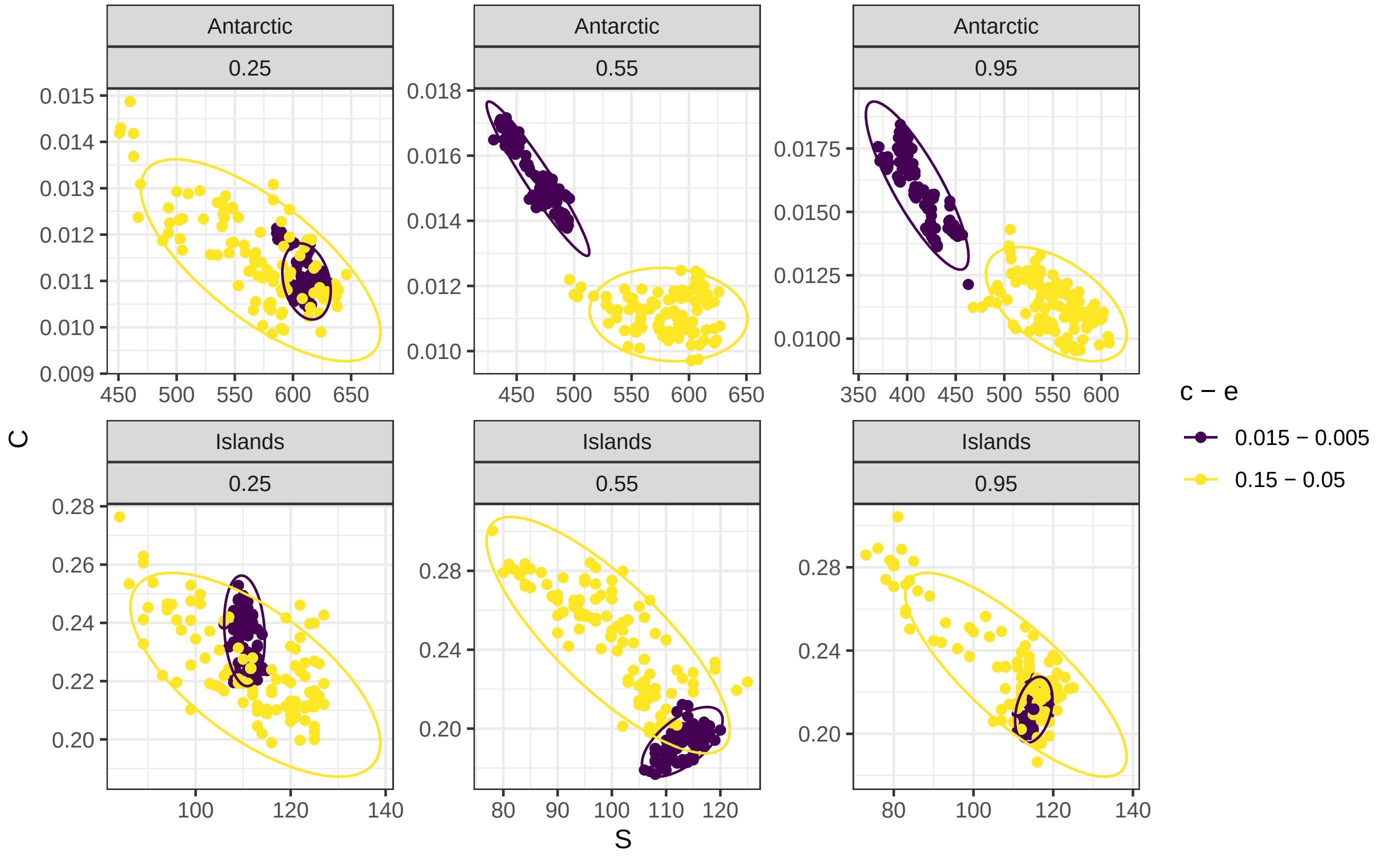


Figure S6: Simulation of the metaweb assembly model keeping $\alpha = c/e$ constant for the Islands and Antarctic metawebs and different combinations of $c$, $e$ and $se$. The points are averages of number of species $S$ and connectance $C$ of the last 100 time steps of the model.
